# Temperature is the key weather determinant of *Aedes albopictus* seasonal activity in southern France

**DOI:** 10.1101/2025.10.10.681559

**Authors:** Paul Taconet, Andrea Radici, Guillaume Lacour, Antoine Mignotte, Pachka Hammami, Didier Fontenille

## Abstract

The presence and the activity of *Aedes albopictus* are a growing concern for nuisance and public health in Europe. Vector control operators, public health officers, and communities look for weather-based decision support systems to inform mosquito management policies. Despite an increasing number of entomological and modelling studies, our incomplete understanding of mosquito population response to weather drivers in natural conditions restricts the development of sound vector management policies.

Here, we aim to clarify the role of weather conditions on *Ae. albopictus* presence and abundance in four sites in southwestern France. We rely on ovitrap longitudinal records collected on a 1-2 weeks basis and weather time series over 2023 and 2024 to model oviposition activity. Our analysis combines a mechanistic model from literature and a new machine learning model fitted on cross-correlated lagged weather predictors.

Both models satisfactorily reproduce the observed oviposition dynamics, correctly predicting the onset and the end of the activity – periods that existing models have often inadequately captured. Temperature plays a major role in triggering the presence of *Ae. albopictus*, explaining the interannual variation of oviposition in all sites, especially in spring and autumn. In fact, warm springs and autumns extend the periods in which *Ae. albopictus* life-history traits (fertility, development, survival) approach their thermal optima. In summer, a more prominent role of rain and humidity emerges among secondary drivers of oviposition intensity.

This work contributes to the development of operational weather-driven forecasting tools for *Ae. albopictus* activity to support vector control operations in different biogeographical contexts.

## Introduction

The Asian tiger mosquito *Aedes albopictus* is an arthropod native to tropical rainforest of Southeast Asia (Paupy et al., 2009). In the last century, facilitated by globalization of shipping, it invaded both tropical and temperate areas worldwide. In mainland France, its first establishment occurred in 2004 in the southeastern regions (Roche et al., 2015). Its biting nuisance and its competence to transmit arbovirus, such as dengue and chikungunya, have raised widespread concerns about its control. In mainland France, the first autochthonous transmission of these arboviruses occurred in 2010 in Nice (dengue) and Fréjus (chikungunya; Franke et al., 2019). Consistently with the intensification of outbreaks in tropical areas, the number of autochthonous cases of *Aedes*-transmitted arboviral disease have gradually increased in Southern Europe, reaching more than 737 cases in mainland France and 353 cases in Italy up to the 8^th^ of October 2025 (Istituto Superiore di Sanità, 2025; Santé Publique France, 2025).

The need to anticipate both mitigate biting nuisance and vector-borne disease risk has prompted vector control operators, public health officials, and citizens to try to interpret the seasonal dynamics of *Ae. albopictus* under the light of possible drivers. These stakeholders adapt their behavior following decision rules – generally supported by scientific evidence – based on a range of models, from simple approaches, as calendar dates (Franke et al., 2019), to complex model ensembles (Da Re et al., 2025). Ultimately, most models assume – either implicitly or explicitly – that the seasonal dynamics of *Ae. albopictus* depend on weather conditions. Eggs need water, usually provided by precipitations, to hatch and develop into larvae and pupae (Paupy et al., 2009). Temperature affects *Ae. albopictus* development, survival and fertility (Mordecai et al., 2019). Humidity is also considered as important for adult survival, and wind patterns plays a minor role on its dispersion (Waldock et al., 2013). Beside these broad principles, there are evidences that *Ae. albopictus* populations exhibit signatures of local adaptation to novel selective pressures, reflecting evolutionary processes that facilitate the establishment in diverse environments. For instance, populations in western Madagascar are adapted to long dry periods (Raharimalala et al., 2012); in temperate areas, they survive to winter thanks to photoperiodically-induced egg diapause, which evolved rapidly during invasion of poleward areas (Lacour et al., 2015; Urbanski et al., 2012), with eggs being resistant to colder temperature compared to their tropical counterparts (Kramer et al., 2021). These adaptive capabilities challenge our possibility to extrapolate current knowledge about *Ae. albopictus* into newly colonized areas, making it more difficult to anticipate adapted measures for the management, surveillance, and control of this species. For example, quantifying the risk that – based on weather conditions – an introduction of an arbovirus-infected person could lead to an outbreak helps operators to prioritize interventions according to available resources. Despite growing research effort to clarify the role of weather determinants of *Ae. albopictus* activity (Brass et al., 2024; Erguler et al., 2016; Kobayashi et al., 2002; Metelmann et al., 2019; Petrić et al., 2021; Tran et al., 2013), local authorities, public health services and mosquito control agencies often lack of solid scientific evidence to support their policies. These are ultimately jeopardized by climate changes, expected to affect the seasonality of this vector (Colón-González et al., 2021).

Our study aims to fill this gap and enable better anticipation of the nuisance and health risk associated with *Ae. albopictus*. We focus on four different ovitrap longitudinal records collected on a 1-2 weeks basis and weather time series over 2023 and 2024 in Occitanie and Nouvelle-Aquitaine, southern France, which have been colonized after 2010. We model oviposition dynamics using both weather-driven statistical and mechanistic approaches, and compare their predictions with ovitrap observations. For these sites, we statistically infer the presence of an interannual trend in terms of ovipositing activity. Therefore, we explore the importance of weather and weather-driven demographical determinants of mosquito trends. Finally, we conclude on some remarks about the possibility of forecasting ovipositing activity and its implications for optimizing mosquito control operations in the context of the climate change.

## Material and methods

### Entomological longitudinal analysis

Our analysis basis on entomological surveillance observations provided by the Altopictus vector-control agency ovitrap collection in four different locations in mainland France, *i.e.* Pérols and Murviel-les-Montpellier (Occitanie), Bayonne and Saint-Médard-en-Jalle (Nouvelle-Aquitaine; Fig. 1). According to the Corine Land Cover classification, the monitored areas correspond to discontinuous urban fabric, whereas some ovitraps in Bayonne were located within continuous urban fabric (European Union’s Copernicus Land Monitoring Service information, 2020). Ovitraps consist of artificial egg-laying containers made of 3-Liter black plastic buckets filled with 2 L of Bti-treated (*Bacillus thuringiensis israelensis* biolarvicide) tap water. A floating polystyrene square (5 x 5 x 2 cm) provided a substrate for oviposition. They are used to monitor the oviposition intensity, *i.e.* the abundance of laid eggs, a proxy of the density of locally active females. These ovitraps have been inspected every 1-2 weeks between 2023 and 2024 (Tab. SI4). Surveillance operations were interrupted from June 2024 onwards in Bayonne and Saint-Médard-en-Jalle. Records have been aggregated using the mean value for each inspection date and site.

**Fig. 1.**
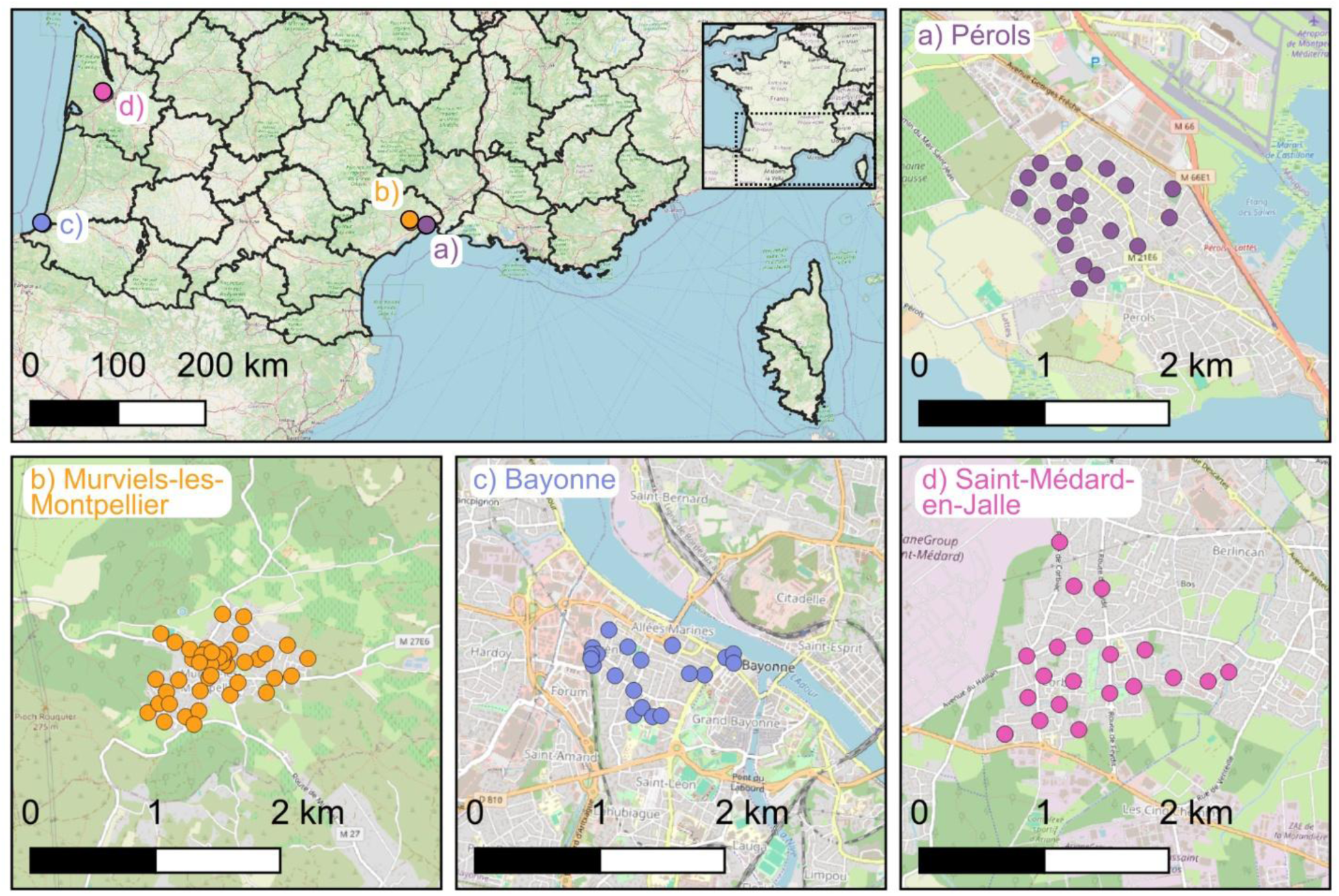
Spatial setup of the Altopictus ovitrap surveillance network in southwestern France: a) Pérols, b) Murviel-les-Montpellier, c) Bayonne, d) Saint-Médard-en-Jalle, over a OpenStreetMap land cover and land use layer.

To assess the impacts of different weather conditions over oviposition activity, we first analyzed the ovitrap series site by site. Each time series was divided into three seasons, *i.e.* “spring” (May-June), “summer” (July-August) and “autumn” (September-October). For each site and season, we determined the presence of an interannual trend via the two-tailed two-samples Wilcoxon test by comparing the statistical distribution of the ovitrap records of the two years of collection.

### Input data

Both mechanistic and statistic models are driven by weather inputs. We retrieved daily precipitation (mm), daily mean, maximum and minimum temperatures (°C), relative humidity (%), average and maximum wind speed (m/s) from MétéoFrance (details in Tab SI4). We retrieved human population density raster at 30”, aggregated over a grid of 0.1° , based on the 2015 GPWv4 data for 2015 (Doxsey-Whitfield et al., 2015).

### Statistical model

In statistical modelling, the response variables and their predictors have to be defined first. Here, we considered the oviposition (measured as “daily number of eggs per ovitrap”), averaged by site and collection date, as the response variable. To distinguish between those weather drivers that induce the onset of mosquito activity and those governing oviposition intensity, we developed two separate models. The first model, called “presence/absence model”, aims to identify the conditions that allow mosquito activity with a binary response (1 = presence of eggs, 0 = absence). The second, called “abundance model”, aims to predict the oviposition intensity as a continuous variable (mean number of eggs per trap per day, restricted to positive trapping events). The modeling pipeline described below was implemented for both model types.

To investigate the temporal relationships between mosquito activity and weather conditions, we first computed cross-correlation maps (CCMs) between the response variable and weekly averaged weather variables at lags ranging from 0 to 12 weeks prior to each trapping date (Curriero et al., 2005). CCMs enable to assess and visualize associations between two time series, such as mosquito abundance and weather conditions, which may be lagged in time. Here, we measured the strength of the association using the Distance correlation coefficient (Székely et al., 2007). Distance correlation captures non-linear associations between variables, hence being more suited to analyze complex ecological systems compared to linear correlation coefficients. These CCMs were computed for each weather variable at each site to identify site-specific patterns, and were also pooled across all sites to assess overall associations for each environmental variable.

For each pooled weather variable, the lag with the strongest correlation to the response variable was retained as a predictor for multivariate modeling. Weather variables displaying weak associations (distance correlation coefficient < 0.1) were discarded. To reduce multicollinearity, we further filtered correlated predictors based on pairwise Pearson correlation (r > 0.7), by selecting those predictors with the highest ecological relevance and interpretability.

The final set of selected predictors was used to train multivariate Random Forest (RF) models (Breiman, 2001). Binary classification RFs were applied for presence modeling, while regression RFs were used for abundance modeling. To avoid overfitting and inflated performance estimates, we implemented a spatial leave-one-site-out cross-validation (LOSO-CV) framework. In this approach, models were iteratively trained on data from three sites and tested on the fourth. This strategy not only mitigates overfitting but also provides a robust assessment of the model’s generalizability to previously unseen locations, for both presence/absence and abundance predictions (Meyer et al., 2018). Finally, the outputs of the presence and abundance models were combined to generate a unified prediction. Specifically, if the predicted probability of presence exceeded the threshold value of 0.5, the corresponding value from the abundance model was retained; otherwise, the prediction was set to zero.

### Mechanistic model

As a mechanistic model, we used the model by Metelmann et al. (2019), currently used and validated in temperate areas in Europe (Barman et al., 2025; Radici et al., 2025), which describes the dynamics of the mosquito population (mosquito/ha) via ordinary differential equations into five stages (eggs *E*, diapausing eggs *E*_*d*_, larvae and pupae as juveniles *J*, unfed immature female adults *I*, mature female adults *A*). These equations specify mortality, development and fertility rates, which rely on environmental (photoperiod *P* and human density *H*) and weather (rainfall *R* and temperature *T*) drivers.

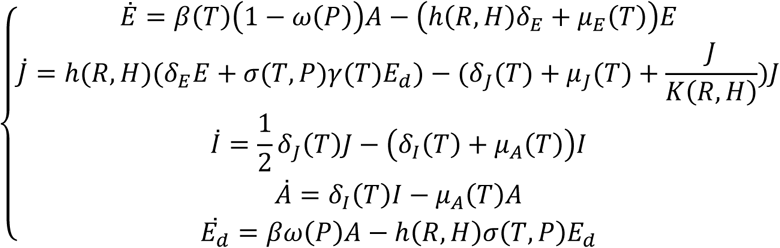

Where *β* represent the fertility rate, *δ*_*E*_, *δ*_*J*_ and *δ*_*I*_the development rate of eggs, juveniles and adults, *μ*_*E*_, *μ*_*J*_, *μ*_*A*_, their mortality, *ω* the fraction of diapausing laid eggs, *h* the hatching rate, *σ* the fraction of diapausing eggs ready to hatch, *γ* the probability of winter survival, *K* the carrying capacity of juveniles (Table SI1 for details on weather dependencies). This model allowed to estimate the observed oviposition abundance via the indicator *LE*_*it*_ of eggs laid per day per hectare in site at time in site *i* at time *t*:

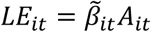

Where *A*_*it*_ is the simulated density of adult female mosquitoes, and *β̃*_*it*_ is the temperature-dependent fertility rate. In order to compare our simulations with observed ovitrap data, we averaged *LE*_*it*_ with a moving window of two weeks and we normalized it with respect to the highest value.

### Model evaluation

We assessed the ability of both models to capture the temporal dynamics of mosquito oviposition by computing the Spearman and Pearson correlation coefficients between predicted and observed egg counts. The statistical significance of each correlation was evaluated using a two-sided Student’s *t*-test.

### Analysis of weather determinants of oviposition

To investigate the role of weather drivers in shaping oviposition dynamics, we interpreted the trained machine learning (ML) models using Variable Importance Plots (VIPs) and Partial Dependence Plots (PDPs). VIPs measure each variable’s contribution to the model performance, while PDPs visualize its average effect on predictions, accounting for interactions with other predictors. These tools enable to highlight which environmental variables mostly influence mosquito presence or abundance.

To complement these global interpretations with fine-scale ecological insights, we applied the Local Interpretable Model-agnostic Explanations (LIME) method (Ribeiro et al., 2016). LIME approximates the complex ML model locally using a simpler and interpretable surrogate (e.g. a linear model). It provides local contributions of each predictor to individual model predictions, indicating both the direction (positive = increasing, negative = decreasing) and the magnitude of influence relative to other predictors at that specific site and time point.

### Analysis of weather-driven demographic determinants of oviposition

For the sites and seasons in which the Wilcoxon test of ovitrap abundance revealed a significant difference between 2023 and 2024 (p-value < 0.05), we analyzed the statistical significance of the weather-dependent demographic rates between those two years using the same Wilcoxon test. The weather-dependent demographic rates considered here are the fertility (*β*), juvenile development (*δ*_*J*_), immature adult development (*δ*_*I*_), egg survival (*e*^−*μE*^), juvenile survival (*e*^−*μJ*^), adult survival (*e*^−*μA*^), hatching (*h*), carrying capacity (*K*). The carrying capacity was considered as a demographic trait as its role in intraspecific competition makes it proportional to survival rate. By testing the inter-annual variability of those rates, we identified those that varied the most “consistently” with oviposition, and can be considered as determinants of mosquito activity.

Hereinafter, we will use “inconsistent” as a synonym for “significantly sharing the opposite interannual trend as the observed oviposition” and, “consistent” as a synonym for “significantly sharing the same interannual trend as the observed oviposition”.

### Software

All analyses were conducted using open-source software. The R programming language (R Core Team, 2024) served as the primary platform. The Wilcoxon statistical analysis of time series was performed using the ‘stat’ package (version 4.4.1). Correlation analyses were performed with the ‘correlation’ package (0.8.8; Makowski et al., 2020). Random forest models were fitted using the ‘caret’ (7.0-1; Kuhn, 2008) and ‘ranger’ (0.17.0) packages, while spatial folds for cross-validation were generated with the ‘CAST’ package (1.0.3). Model interpretability analyses relied on the ‘vip’ (0.4.1; Greenwell & Boehmke, 2020) and ‘pdp’ (‘0.8.2’; Greenwell, 2017) packages to produce variable importance and partial dependence plots, respectively, and the ‘lime’ package (0.5.3, Hvitfeldt et al., 2022) to perform the LIME analysis. Among the inputs of the mechanistic model, we retrieved the photoperiod (in hours) using the ‘suncalc’ (0.5.1), based on the latitude, longitude and date.

## Results

### Analysis of oviposition and weather time series

Compared to 2023, the oviposition onset of 2024 occurred later in the season for each site except Bayonne (Fig. 2, Table SI4). For both Pérols and Murviel-les-Montpellier, the last positive ovitrap was found later in the year in 2023 compared to 2024. Despite the variability of ovitraps records, the Wilcoxon test revealed the presence of a significant difference in observed oviposition abundance between seasons (Fig. 2, Fig. 6). In Pérols, oviposition followed a bimodal pattern in 2023 and was higher in spring and autumn (but not significantly), contrarily to summer season, where oviposition was higher in 2024 (Fig. 2a, Fig. 6a). In Murviel-les-Montpellier, oviposition was always higher in 2023, but this difference was not significant in summer (Fig. 2b, Fig. 6b). In Saint-Médard-en-Jalle and Bayonne, spring oviposition was higher in 2023 than in 2024 (Fig. 2c,d, Fig. 6c,d). Every season in every site was warmer in 2023 compared to 2024 (with the exception of summer in Bayonne and Saint-Médard-en-Jalle, in which temperatures remained roughly stable). Except for Pérols, where rainfall was higher in every season in 2024 compared to 2023, there was no clear interannual trend in precipitation (Tab. SI4).

**Fig. 2.**
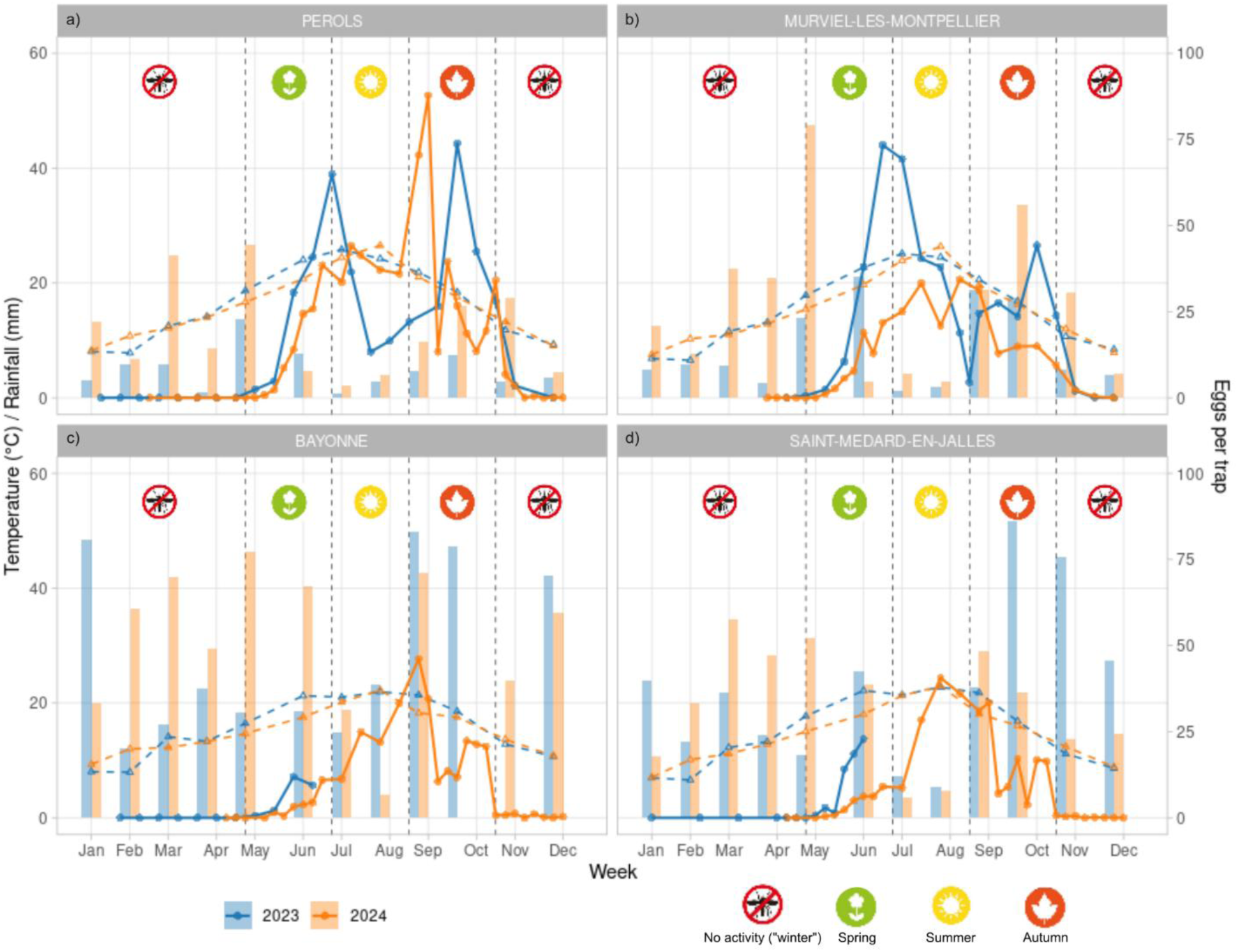
Ovitrap data (solid line; mean value for each reading) in 2023 - 2024 in the 4 sites, together with average daily temperature (dashed line; averaged with a monthly moving window) and monthly rainfall (bars).

### Lagged associations between weather and oviposition dynamics

Cross-correlation analyses between mosquito dynamics and weather variables revealed distinct temporal patterns for eggs presence and abundance across the study sites (Fig.s SI2 and SI3). Temperature-related variables – average, minimum, and maximum temperature – consistently emerged as the most correlated predictors for both presence and abundance. For presence, the strongest associations are generally observed at 1-8 weeks lags, while for abundance, they are observed at shorter time lags (1-4 weeks). Relative humidity and rainfall-related variables displayed weaker and more heterogeneous correlations, with peak correlations generally occurring at longer time lags (up to 12 weeks). Wind-related variables generally exhibited the weakest associations with mosquito dynamics, with few exceptions. The temporal structure of associations displayed a notable degree of consistency across sites, especially for temperature.

Among the weather variables, for the presence/absence RF model, we selected: (i) average temperature between the 1^st^ and 9^th^ week (preceding oviposition; TM_0_8), and (ii) relative humidity from the 6^th^ to the 12^th^ week (UM_5_11). For the abundance RF model, the retained predictors included: (i) average temperature from the 1^st^ to the 5^th^ week (TM_0_4), (ii) relative humidity from the 1^st^ to the 12^th^ week (UM_0_11), and (iii) cumulative rainfall from the 2^nd^ to the 6^th^ week (RR_1_5).

### Model performance assessment

Both statistic and mechanistic models globally captured the seasonal trend (Fig. 3) – except ML models in Bayonne (Fig. 3g). The beginning and the end of the seasons were globally better captured by ML models, especially in those locations where the whole time series was available (Pérols, Murviel; Fig. 3e, f). In the same locations, ML models were particularly good at capturing the inter-seasonal trends (peaks and downs). The predictions of the models were always significantly close to the observations (Tab. 1).

**Fig. 3.**
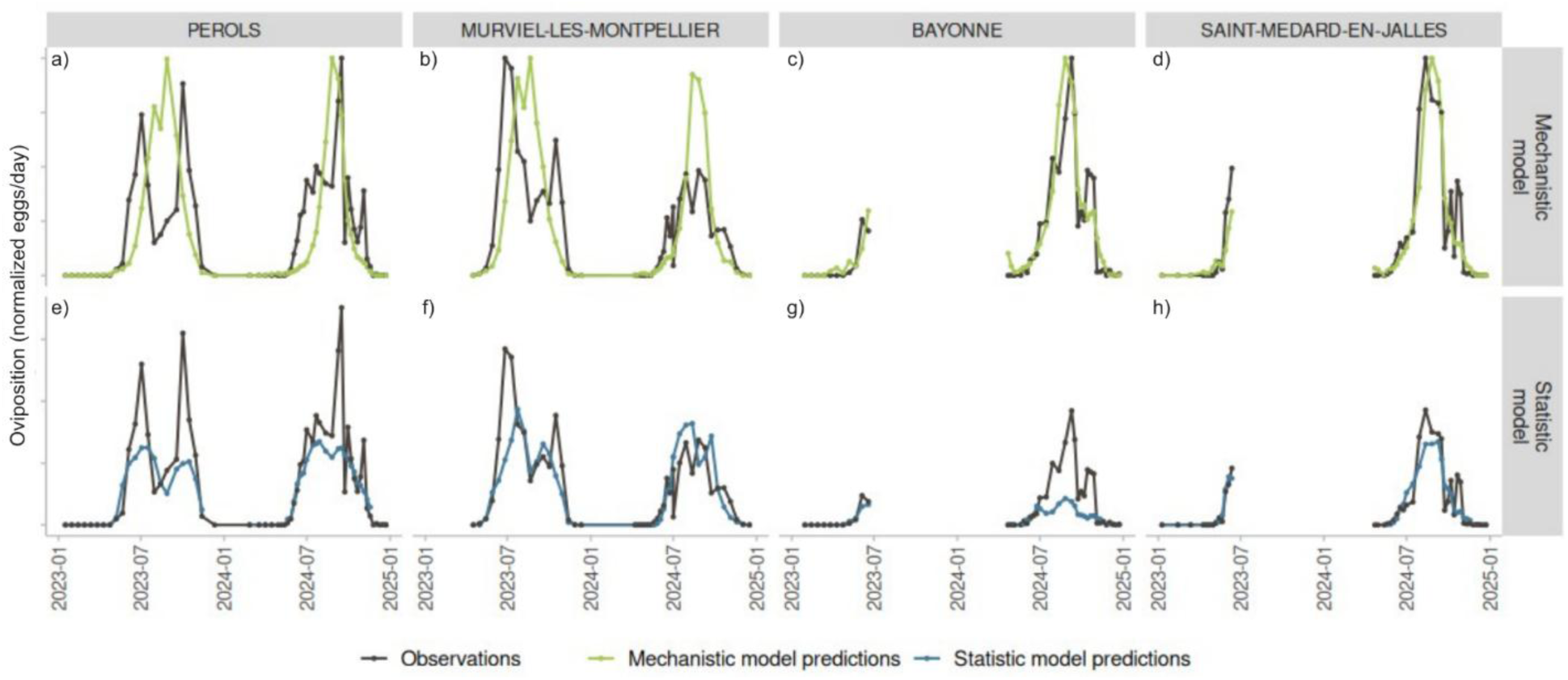
Observed oviposition vs simulated oviposition in each site. First row: normalized observations versus predictions of the mechanistic model. Second row: absolute observations versus the machine-learning model predictions.

**Table 1.**
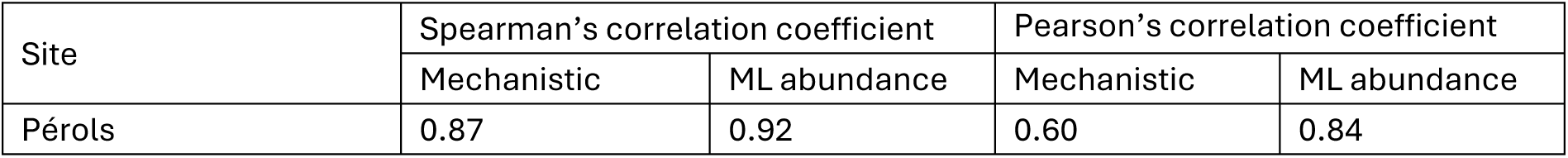
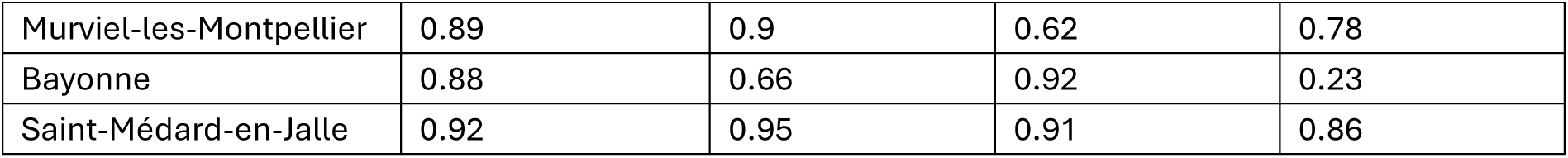
Performances of the abundance models. All the correlation values are significative at a p-value < 0.001.

### Analysis of environmental determinants

#### Pooled-site effects of weather variables on oviposition activity

Variable importance plots (VIPs) and partial dependence plots (PDPs) highlighted the dominant role of temperature in predicting both mosquito presence and abundance (Fig. 4). For the presence model, average temperature between the 1^st^ and 9^th^ weeks (before oviposition; TM_0_8) was by far the most important predictor across all sites, while relative humidity during weeks 6 to 12 (UM_5_11) had only a minor impact (Fig. 4a). The PDPs showed a sharp increase in the probability of mosquito presence as TM_0_8 rises from approximately 11°C to 18 °C, after which the probability saturated (Fig. 4b). Relative humidity exhibited a weaker and more complex relationship – generally negative – with presence probability (Fig. 4c).

**Fig. 4.**
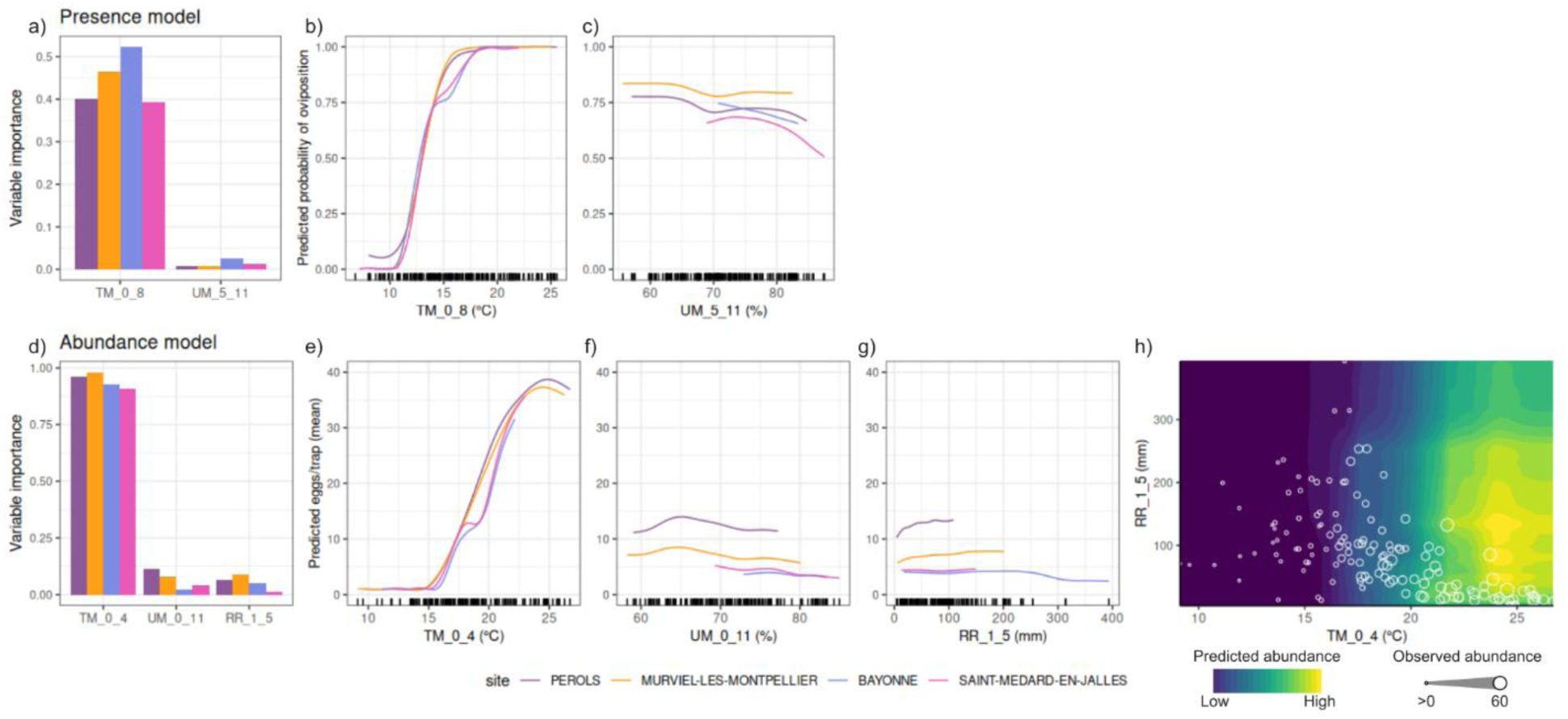
VIPs and PDPs for the mosquito presence (top) and abundance (bottom) models. Bar plots (a,d; left) show variable importance, indicating the relative contribution of each environmental predictor to the RF model. Line plots (b, c, e, f, g; right) display partial dependance, representing the marginal effect of each predictor on the response variable while averaging over the influence of other variables. The bivariate PDP in the bottom panel (h) illustrates the combined effect of temperature (TM_0_4) and rainfall (RR_1_5). For the presence model, the response variable is the probability of oviposition occurrence; for the abundance model, it is the predicted number of eggs per trap (restricted to presence-only data). These plots are based on models trained on pooled data across all sites.

In the abundance model, average temperature weeks 1 to 5 (TM_0_4) was the most influential predictor, followed by relative humidity (UM_0_11) and cumulative rainfall (RR_1_5; Fig. 4d). The PDPs revealed a strong, linear increase in predicted egg abundance with TM_0_4 ranging from 15 °C to ∼24 °C, after which the abundance saturated and then decreased (Fig. 4e). In contrast, higher relative humidity was generally associated with a decrease in predicted abundance (Fig. 4f). The effects of rainfall differed across sites, showing a slightly positive effect on oviposition intensity in Pérols and Murviel-lès-Montpellier, while appearing negligible in Bayonne and Saint-Médard-en-Jalles (Fig. 4g). The pooled bivariate PDP for TM_0_4 and RR_1_5 highlighted an optimal range of conditions for oviposition activity, with maximum predicted egg abundance occurring near 25 °C average temperature of the previous 5 weeks (TM_0_4) and approximately 150 mm cumulative rainfall over the 2^nd^ to 5^th^ weeks before deposition (RR_1_5; Fig. 4h).

#### Local spatio-temporal effects of weather variables on oviposition activity

Across all sites and weeks, temperature consistently emerged as the most influential predictor driver, with strong positive contributions aligning closely with observed peaks in egg abundance, particularly during the warmer months (Fig. 5, and SI4-7).

**Fig. 5.**
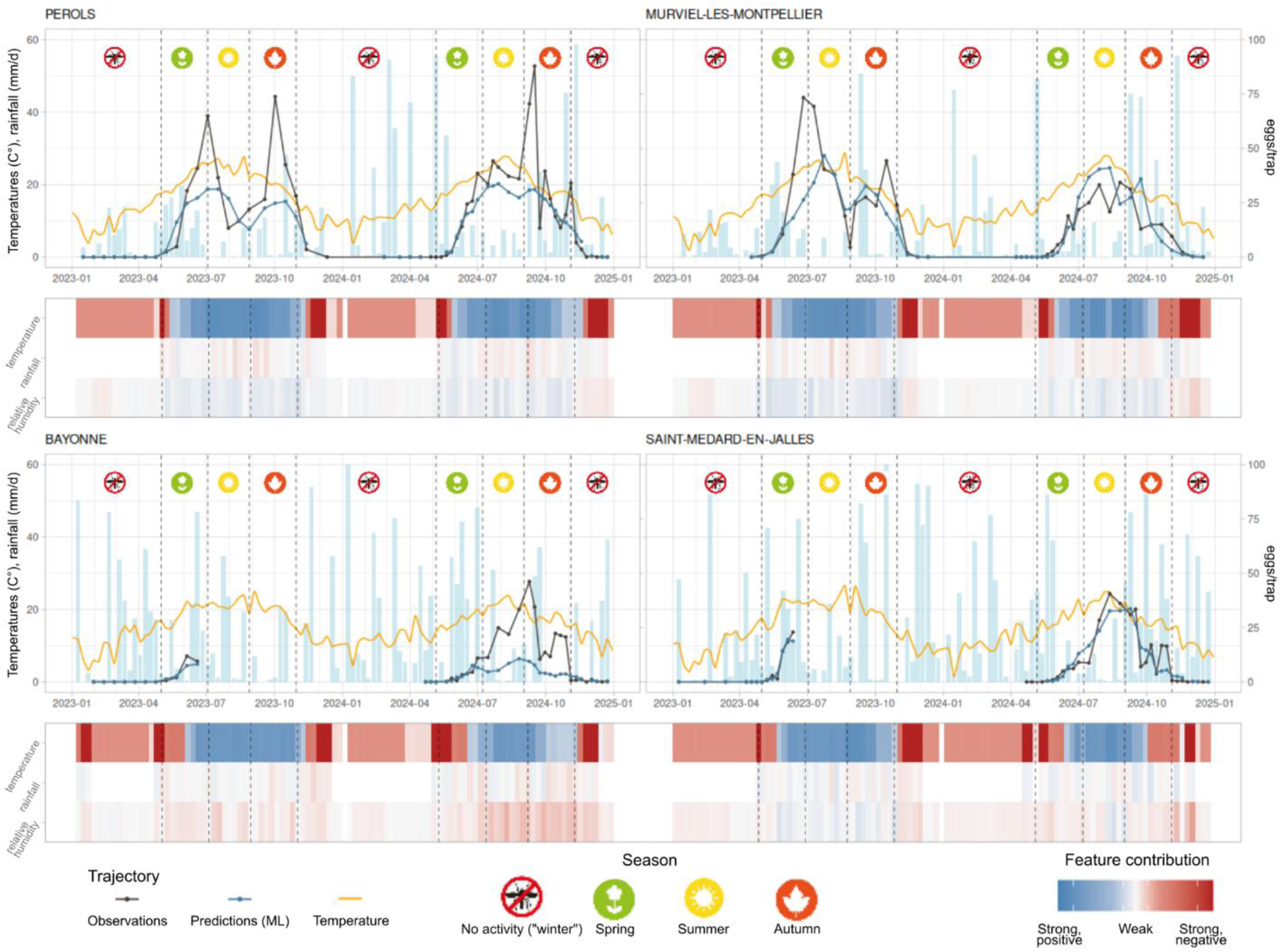
Local contribution of environmental predictors to predicted mosquito abundance using LIME. Each colored tile represents the contribution of a given environmental variable to the model’s predicted egg presence/ and abundance for a specific site and time point (weekly resolution, 2023–2024). Colors indicate whether the variable contributed positively (blue) or negatively (red) to the prediction, with color intensity proportional to the magnitude of the effect.

**Fig. 6.**
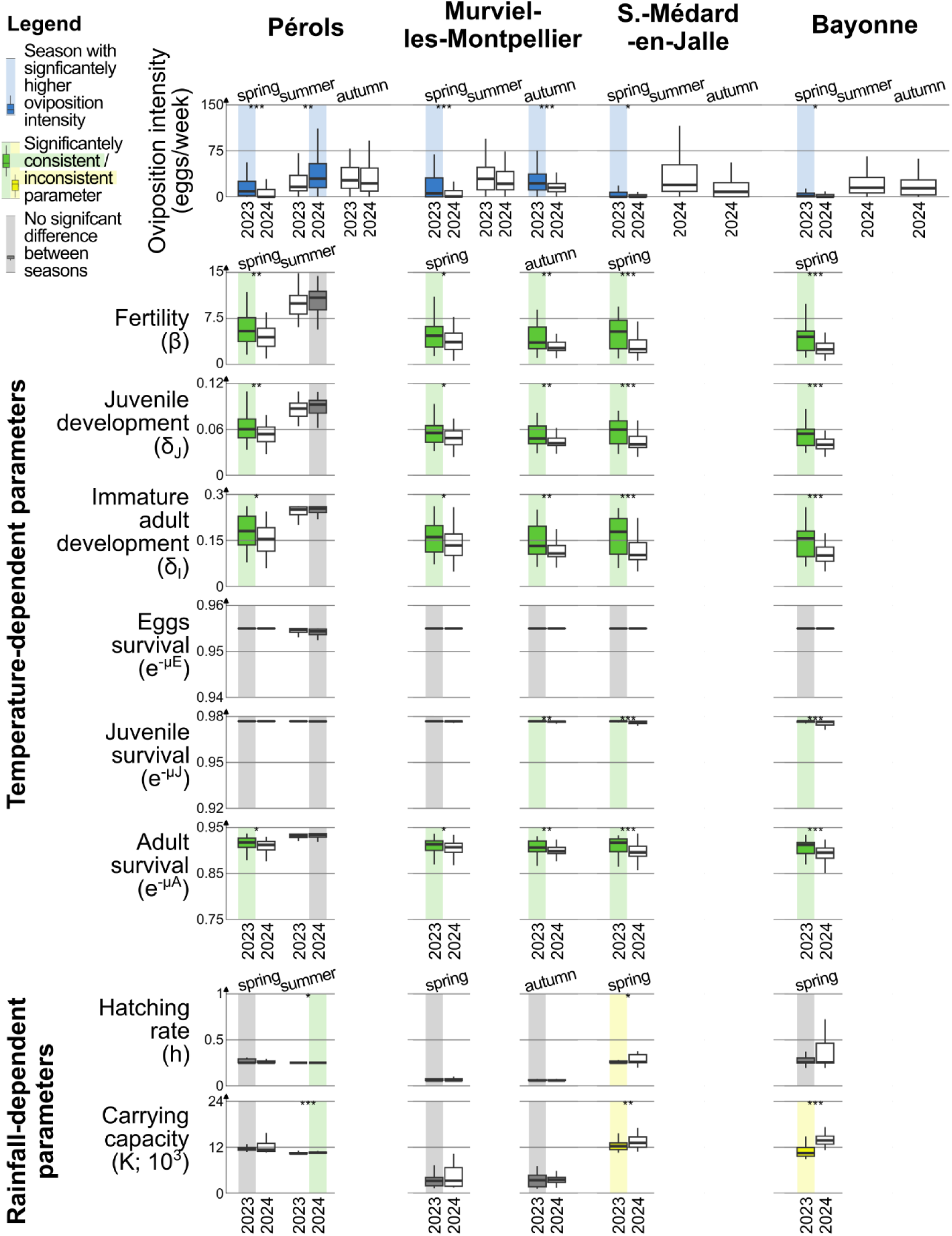
Consistency analysis per site, year and season between oviposition intensity (boxplots in the first row) against mechanistic weather-driven parameters (boxplots from the second row). For each site and season, the boxplots corresponding to the year of highest oviposition are colored in blue. Then, the boxplots of those combinations of site and season in which the parameters varied consistently(/inconsistently) with oviposition are colored in green(/yellow). Gray boxplots mean lack of significance. The stars indicate the p-value (*: < 0.05, ***: < 0.01, ***: < 0.001) according to the one-to-one Wilcoxon test.

Rainfall and humidity, by contrast, showed more variable and context-dependent effects. Notably, in Pérols and Murviel-lès-Montpellier, distinct declines in predicted abundance were observed during summer (Fig. 5a,b). In this season, temperature remained positively associated with oviposition intensity, whereas rainfall made a negative contribution to the model predictions.

### Analysis of weather-driven mosquito demographic determinants

In the mechanistic model, temperature-related parameters, with the exclusion of egg survival, were globally consistent with ovitrap observations (Fig. 6). Fertility (*β*), juvenile development (*δ*_*J*_), immature adult development (*δ*_*I*_) and adult survival (*e*^−*μA*^) were consistent 5 times out of 6, juvenile survival (*e*^−*μJ*^) 3 out of 6. In all the other cases, the trend was neither significant nor inconsistent. By contrast, observed oviposition was consistent with rainfall-dependent demographic traits – hatching (*h*) and carrying capacity (*K*) – only in summer in Pérols. In all other cases, they were either not significant or inconsistent.

## Discussion

### Average temperatures of the previous weeks are the main drivers of oviposition in Southern France

In the study area, interannual trends of ovipositing activity – a widespread proxy used to monitor the activity of adult females (Da Re et al., 2024) having taken a blood meal to develop their eggs – can be easily explained by a limited number of weather drivers, which impact different life-history traits. Average temperature of the previous 9 weeks is by far the main determinant of the presence of eggs, and of the previous 5 for their abundance. The importance of considering lagged temperatures into the description of oviposition dynamics in southern Europe is confirmed by recent research – despite the disagreement with these studies on the amplitude of the lag. For instance, Torina et al. (2023), found that average temperature of the previous 5 weeks is the most correlated driver to observed oviposition intensity in Palermo, Italy, while Da Re et al., (2025) found that average temperature of the third-to-last week is the most important determinant of oviposition intensity over the area covered by the VectAbundance survey in southeastern Europe (Da Re et al., 2024). These differences may reflect either local population dynamics or the results of different modelling choices. Concerning diapause end, it is interesting to notice that our results allow to correctly capture and satisfactorily explain the onset of mosquito activity also neglecting the photoperiod, the main driver of egg diapause (i.e., the population overwintering through the production of eggs that will hatch in spring; Lacour et al., 2015; Lounibos et al., 2003; Sturiale & Armbruster, 2023).

### Warm temperatures facilitate longer and more intense activity seasons of *Aedes albopictus* in a temperate climate

Wide research focuses on temperature as the main driver of the ecology of ectotherm populations. Mordecai et al. (2019) formalized the performance of a corpus of mosquito life-history traits as an (asymmetric) bell-shaped function of temperature (Briere et al., 1999; Waldock et al., 2013). The optimal thermal response of *Ae. albopictus* on several key biological traits varies between 24.2 °C (survival of aquatic stages) to 32.6 °C (adult development rate; de Souza & Weaver, 2024). For temperatures lower than the optimal ones, as it regularly occurs in temperate countries, a temperature increase corresponds to an increment of mosquito fitness (de Souza & Weaver, 2024). Our analysis of demographic determinants in the mechanistic model corroborates this idea; temperature-dependent traits (i.e. survival, development or fertility rates) varied consistently with oviposition intensity across all sites during cool seasons (spring and autumn) characterized by suboptimal temperatures.

It is worth noticing that climate change increases the likelihood of warmer springs and autumns, with important implications on the epidemiological risk. In fact, in a temperate climate such as that of southern France, the increase of temperatures could lead to the extension of the season at risk of autochthonous transmission of arboviruses, as noted by other studies (Colón-González et al., 2021; Radici et al., 2025). In fact, temperature affects the extrinsic incubation time of arboviruses, which develop and are transmitted more easily at hot temperatures (Liu-Helmersson et al., 2014). It is therefore advisable that the calendar for enhanced surveillance, which currently begins in May in mainland France (Franke et al., 2019), should be adjusted accordingly.

### Rainfall has a minor role in sustaining mosquito activity in southern France

The analysis of environmental determinants indicates that rainfall plays a secondary role in shaping population abundance dynamics, while it does not affect the presence. Consistently, the analysis of environmentally-driven demographics suggests that rainfall becomes determinant only during summer. The primary effects of rainfall are to trigger egg hatching and to increase the availability of larval breeding sites, the main limiting resource to mosquito population growth (Waldock et al., 2013). Yet, the importance of rainfall in sustaining larval survival – and therefore mosquito population – in urban settings is increasingly debated (Boyer et al., 2014). Urbanization has led to an increased capacity to retain water – whether from rainfall or irrigation – due to the expansion of impervious surfaces (gutter silt traps, raised terraces, roadworks, flowers pots, rainfall collectors, etc.). In many cases, human irrigation compensates for reduced precipitation, providing persistent larval refugia that reduce mosquito populations dependence from natural water supply (Li et al., 2014). It is not surprising that rainfall-dependent parameters only become the limiting factors to oviposition only during the driest season in the driest locations (∼ 21 mm of cumulative rainfall in June and July in Pérols, while Bayonne has ∼110; Tab. SI4). Moreover, Fonseca et al. (2015) suggested that in autumn, females laying diapausing eggs exhibit a preference for larger containers; this behavioral shift reduces the dependance of the first generations of larvae on rainfall abundance (or scarce evaporation).

The VIP analysis highlights a secondary role of relative humidity, which affects negatively both presence and abundance of oviposition. Relative humidity has long been recognized as a challenging variable to embed into a modelling scheme due to its complex interactions with other weather variables (Waldock et al., 2013). For instance, high humidity is expected to enhance eggs survival at 26 °C, but the same does not hold consistently across other temperatures (Juliano et al., 2002). The effect of humidity on adult mosquitoes is even less understood: European populations of Asian tiger mosquito have been found also in regions with relative humidity levels below 35% (Waldock et al., 2013). In conclusion, the range of values we measured in the study area is probably not broad enough – between 60 and 85% – to have an appreciable effect on mosquito dynamics.

### The effects of extreme weather events on the mosquito activity in southern France

Summer temperatures may exceed the optimal range of *Ae. albopictus*. This occurred in the warmer sites of Pérols and Murviel-les-Montpellier, where the PDP reveal a slight decrease of abundance for average monthly temperatures exceeding 24 °C. While on-field evidence of this phenomenon in temperate climate is poor, other modelling study document the negative effect of summer heatwaves on mosquito fitness (Garrido Zornoza et al., 2024). As a matter of fact, it is difficult to disentangle the effect of heatwaves from that of droughts, as in temperate climate they often occur simultaneously. Notably, in Pérols and Murviel-lès-Montpellier, distinct declines in predicted abundance were observed during summer 2023, despite suboptimal but still favorable temperature conditions. In these dates, the LIME analysis provided an insight on the role of low rainfall, which makes a negative contribution to the model predictions, suggesting that low or insufficient precipitation results in limited oviposition. In contrast, in Bayonne and Saint-Médard-en-Jalles, rainfall and humidity show minimal or inconsistent influence across the season, indicating a reduced predictive value of these variables under the prevailing local oceanic conditions.

Across the available time series, we have not observed extreme rainfall events. Several studies reported that excessive rain may flush out breeding sites, potentially disrupting mosquito population dynamics (Waldock et al., 2013). However, the 2014 epidemic of chikungunya in the Montpellier area – which includes Pérols and Murviel-les-Montpellier – following an exceptional rain event (299 mm on the 29th of September) lead researchers to reconsider the effect of extreme precipitation. In such cases, heavy rainfall may also create an unprecedented number of new breeding sites, ultimately boosting the mosquito population (Roiz et al., 2015; White et al., 2025). While our analysis highlights that rainfall plays a secondary role in driving oviposition intensity, the PDPs indicate that cumulative rainfall exceeding 150–250 mm over the preceding five weeks may be associated with a decrease in oviposition intensity.

### Limitations of the study

Some caveats must be considered before generalizing the outcomes of this study. The results we obtained need to be contextualized within a specific environment in terms of landscape, climate and social features – specifically, a temperate climate within a western

European urban fabric. These highly anthropized sites are located approximately at the same latitude, close to the coasts. We believe that our results can be generalized to other contexts that share similar environmental features in terms of climate and urbanization. However, extending entomological observations both in time and space would help capturing a wider range of environmental conditions and associated mosquito responses. This would help solving site-specific discrepancies (such as underestimating extreme abundance peaks), typical of ML modelling, ultimately strengthening its operational value for vector control.

Some obstacles to the generalizability of any study on *Ae. albopictus* are its high phenotypical plasticity and adaptation capability, for example of diapause, whose rapid evolution due to the latitudinal expansion of this species requires constant surveillance. Although photoperiod has a minor role in our study, it remains the primary regulator of winter egg diapause, and its evolution would alter this species’ seasonality. Some studies suggested that year-round activity may become possible in the future due to climate change in Southern Europe (Del Lesto et al., 2022).

### Weather-driven models as decision support systems in mosquito management

Our models accurately predicted the onset of mosquito activity after diapause, a hard benchmark for many entomological models, as well as variations of abundances during the mosquito activity season. These results reinforce the complementary of statistical and mechanistic tools to analyze vector dynamics – as previously noted by Tran et al. (2020) in La Réunion – but also their potential to support operational interventions. The outcome of these models can assist vector control operators in prioritizing periods or geographical areas for vector control interventions, or enable health authorities to activate targeted surveillance activities. For instance, the early prediction of the spring onset of mosquito activity would allow for the planning of timing anti-larval interventions. Similarly, health services and mosquito control operators may be mobilized as average temperatures exceed ∼18°C over five weeks, at which we obtained the steepest abundance increase, which has consequences over arbovirus transmission. The integration of such modelling outcomes into the routines of mosquito control operators, health services, and local authorities would allow quick actions to abate the health risk.

While examples of similar operational tools exist – such as the ARBOCARTO framework (Marti et al., 2022), which is already used in some regions of France and abroad, or the early warning system described by Díaz et al. (2024) and used in the Caribbean – there remains substantial room to improve their societal uptake. The development of ARBOCARTO highlights that creating effective tools to predict mosquito activity requires strong coordination among a wide range of stakeholders, which is not limited to the collaboration between field entomologists and ecological modelers. Beyond the scientific community, such an effort calls for collaboration between public health authorities, vector-control operators, local political officials, urban management services, and representatives of civil society. Only through this integrative, multi-actor approach can surveillance tools be effectively translated into actionable strategies for public health and vector control. Lastly, to enhance the effectiveness of these models and tools, they should be designed to forecast mosquito activity (in the future) – and not only predict it (in the present).

## Acknowledgements

The authors acknowledge the support of RIVOC funded by the Occitanie region in France for the VECTOCLIM project. We thanks Colombine Bartholomée, Annelise Tran, Renaud Marti, Florence Fournet and Cyril Caminade for fruitful exchanges.

## References

Barman, S., Semenza, J. C., Singh, P., Sjödin, H., Rocklöv, J., & Wallin, J. (2025). A climate and population dependent diffusion model forecasts the spread of Aedes Albopictus mosquitoes in Europe. Communications Earth & Environment, 6(1), 276. 10.1038/s43247-025-02199-z

Boyer, S., Foray, C., & Dehecq, J.-S. (2014). Spatial and Temporal Heterogeneities of Aedes albopictus Density in La Reunion Island: Rise and Weakness of Entomological Indices. PLOS ONE, 9(3), e91170. 10.1371/journal.pone.0091170

Brass, D. P., Cobbold, C. A., Purse, B. V., Ewing, D. A., Callaghan, A., & White, S. M. (2024). Role of vector phenotypic plasticity in disease transmission as illustrated by the spread of dengue virus by Aedes albopictus. Nature Communications, 15(1), 7823. 10.1038/s41467-024-52144-5

Breiman, L. (2001). Random Forests. Machine Learning, 45(1), 5–32. 10.1023/A:1010933404324

Briere, J.-F., Pracros, P., Le Roux, A.-Y., & Pierre, J.-S. (1999). A Novel Rate Model of Temperature-Dependent Development for Arthropods. Environmental Entomology, 28(1), 22–29. 10.1093/ee/28.1.22

Colón-González, F. J., Sewe, M. O., Tompkins, A. M., Sjödin, H., Casallas, A., Rocklöv, J., Caminade, C., & Lowe, R. (2021). Projecting the risk of mosquito-borne diseases in a warmer and more populated world: A multi-model, multi-scenario intercomparison modelling study. The Lancet Planetary Health, 5(7), e404–e414. 10.1016/S2542-5196(21)00132-7

Curriero, F., Shone, S., & Glass, G. (2005). Cross Correlation Maps: A Tool for Visualizing and Modeling Time Lagged Associations. Counselling Psychology Review, 5(3), 267–275. 10.53841/bpscpr.2016.31.1.67

Da Re, D., Marini, G., Bonannella, C., Laurini, F., Manica, M., Anicic, N., Albieri, A., Angelini, P., Arnoldi, D., Bertola, F., Caputo, B., De Liberato, C., della Torre, A., Flacio, E., Franceschini, A., Gradoni, F., Kadriaj, P., Lencioni, V., Del Lesto, I., … Rosà, R. (2025). Modelling the seasonal dynamics of Aedes albopictus populations using a spatio-temporal stacked machine learning model. Scientific Reports, 15(1), 3750. 10.1038/s41598-025-87554-y

Da Re, D., Marini, G., Bonannella, C., Laurini, F., Manica, M., Anicic, N., Albieri, A., Angelini, P., Arnoldi, D., Blaha, M., Bertola, F., Caputo, B., De Liberato, C., della Torre, A., Flacio, E., Franceschini, A., Gradoni, F., Kadriaj, P., Lencioni, V., … Rosà, R. (2024). VectAbundance: A spatio-temporal database of Aedes mosquitoes observations. Scientific Data, 11(1), 636. 10.1038/s41597-024-03482-y

de Souza, W. M., & Weaver, S. C. (2024). Effects of climate change and human activities on vector-borne diseases. Nature Reviews Microbiology, 1–16. 10.1038/s41579-024-01026-0

Del Lesto, I., De Liberato, C., Casini, R., Magliano, A., Ermenegildi, A., & Romiti, F. (2022). Is Asian tiger mosquito (Aedes albopictus) going to become homodynamic in Southern Europe in the next decades due to climate change? Royal Society Open Science, 9(12), 220967. 10.1098/rsos.220967

Díaz, A. R., Rollock, L., Boodram, L.-L. G., Mahon, R., Best, S., Trotman, A., Meerbeeck, C. J. V., Fletcher, C., Dunbar, W., Lippi, C. A., Lührsen, D., Sorensen, C., Muñoz, Á. G., Ryan, S. J., Stewart-Ibarra, A. M., & Lowe, R. (2024). A demand-driven climate services for health implementation framework: A case study for climate-sensitive diseases in Caribbean Small Island Developing States. PLOS Climate, 3(10), e0000282. 10.1371/journal.pclm.0000282

Doxsey-Whitfield, E., MacManus, K., Adamo, S. B., Pistolesi, L., Squires, J., Borkovska, O., & Baptista, S. R. (2015). Taking Advantage of the Improved Availability of Census Data: A First Look at the Gridded Population of the World, Version 4. 226–234.

Erguler, K., Smith-Unna, S. E., Waldock, J., Proestos, Y., Christophides, G. K., Lelieveld, J., & Parham, P. E. (2016). Large-scale modelling of the environmentally-driven population dynamics of temperate aedes albopictus (Skuse). PLoS ONE, 11(2), 1–28. 10.1371/journal.pone.0149282

European Union’s Copernicus Land Monitoring Service information. (2020). CORINE Land Cover 2018 (vector/raster 100 m), Europe, 6-yearlye [Dataset]. 10.2909/960998c1-1870-4e82-8051-6485205ebbac

Fonseca, D. M., Kaplan, L. R., Heiry, R. A., & Strickman, D. (2015). Density-Dependent Oviposition by Female Aedes albopictus (Diptera: Culicidae) Spreads Eggs Among Containers During the Summer but Accumulates Them in the Fall. Journal of Medical Entomology, 52(4), 705–712. 10.1093/jme/tjv060

Franke, F., Giron, S., Cochet, A., Jeannin, C., Leparc-Goffart, I., De Valk, H., & others. (2019). Émergences de dengue et de chikungunya en France métropolitaine, 2010–2018. Bull. Epidémiol. Hebd, 19(20), 374–382.

Garrido Zornoza, M., Caminade, C., & Tompkins, A. M. (2024). The effect of climate change and temperature extremes on Aedes albopictus populations: A regional case study for Italy. Journal of The Royal Society Interface, 21(220), 20240319. 10.1098/rsif.2024.0319

Greenwell, B. M. (2017). pdp: An R Package for Constructing Partial Dependence Plots. The R Journal, 9(1), 421–436.

Greenwell, B. M., & Boehmke, B. C. (2020). Variable Importance Plots—An Introduction to the vip Package. The R Journal, 12(1), 343–366.

Hvitfeldt, E., Lin Pedersen, T., & Benesty, M. (2022). lime: Local Interpretable Model-Agnostic Explanations — lime-package. https://lime.data-imaginist.com/reference/lime-package.html

Istituto Superiore di Sanità. (2025). Arbovirosi • bollettini periodici arbovirosi. https://www.epicentro.iss.it/arbovirosi/dashboard

Juliano, S. A., O’Meara, G. F., Morrill, J. R., & Cutwa, M. M. (2002). Desiccation and thermal tolerance of eggs and the coexistence of competing mosquitoes. Oecologia, 130(3), 458–469. 10.1007/s004420100811

Kobayashi, M., Nihei, N., & Kurihaha, T. (2002). Analysis of northern distribution of Aedes albopictus (Diptera: Culicidae) in Japan by geographical information system. Journal of Medical Entomology, 39(1), 4–11. 10.1603/0022-2585-39.1.4

Kramer, I. M., Pfeiffer, M., Steffens, O., Schneider, F., Gerger, V., Phuyal, P., Braun, M., Magdeburg, A., Ahrens, B., Groneberg, D. A., Kuch, U., Dhimal, M., & Müller, R. (2021). The ecophysiological plasticity of *Aedes aegypti* and *Aedes albopictus* concerning overwintering in cooler ecoregions is driven by local climate and acclimation capacity. Science of The Total Environment, 778, 146128. 10.1016/j.scitotenv.2021.146128

Kuhn, M. (2008). Building Predictive Models in R Using the caret Package. Journal of Statistical Software, 28, 1–26. 10.18637/jss.v028.i05

Lacour, G., Chanaud, L., L’Ambert, G., & Hance, T. (2015). Seasonal Synchronization of Diapause Phases in Aedes albopictus (Diptera: Culicidae). PLoS ONE, 10(12), 1–16. 10.1371/journal.pone.0145311

Li, Y., Kamara, F., Zhou, G., Puthiyakunnon, S., Li, C., Liu, Y., Zhou, Y., Yao, L., Yan, G., & Chen, X.-G. (2014). Urbanization Increases Aedes albopictus Larval Habitats and Accelerates Mosquito Development and Survivorship. PLOS Neglected Tropical Diseases, 8(11), e3301. 10.1371/journal.pntd.0003301

Liu-Helmersson, J., Stenlund, H., Wilder-Smith, A., & Rocklöv, J. (2014). Vectorial capacity of Aedes aegypti: Effects of temperature and implications for global dengue epidemic potential. PloS One, 9(3), e89783. 10.1371/journal.pone.0089783

Lounibos, L. P., Escher, R. L., & Lourenço-De-Oliveira, R. (2003). Asymmetric Evolution of Photoperiodic Diapause in Temperate and Tropical Invasive Populations of Aedes albopictus (Diptera: Culicidae). Annals of the Entomological Society of America, 96(4), 512–518. 10.1603/0013-8746(2003)096%255B0512:AEOPDI%255D2.0.CO;2

Makowski, D., Ben-Shachar, M. S., Patil, I., & Lüdecke, D. (2020). Methods and Algorithms for Correlation Analysis in R. Journal of Open Source Software, 5(51), 2306. 10.21105/joss.02306

Marti, R., Castets, M., Demarchi, M., Catry, T., Besnard, G., Chouin, S., Clément, C., Esteve-Moussion, I., Etienne, M., Foussadier, R., Habchi-Hanriot, N., Jouanthoua, F., L’Ambert, G., Godal, A., & Tran, A. (2022). ARBOCARTO: An operational spatial modeling tool to predict the dynamics of Aedes mosquito species from weather and environmental variables [Conference_item]. ESA. https://agritrop.cirad.fr/602160/

Mcmaster, G. S., & Wilhelm, W. W. (1997). Growing degree-days: One equation, two interpretations. Agricultural and Forest Meteorology, 87(1).

Metelmann, S., Caminade, C., Jones, A. E., Medlock, J. M., Baylis, M., & Morse, A. P. (2019). The UK’s suitability for Aedes albopictus in current and future climates. Journal of the Royal Society Interface, 16(152). 10.1098/rsif.2018.0761

Meyer, H., Reudenbach, C., Hengl, T., Katurji, M., & Nauss, T. (2018). Improving performance of spatio-temporal machine learning models using forward feature selection and target-oriented validation. Environmental Modelling & Software, 101, 1–9. 10.1016/j.envsoft.2017.12.001

Mordecai, E. A., Caldwell, J. M., Grossman, M. K., Lippi, C. A., Johnson, L. R., Neira, M., Rohr, J. R., Ryan, S. J., Savage, V., Shocket, M. S., Sippy, R., Stewart Ibarra, A. M., Thomas, M. B., & Villena, O. (2019). Thermal biology of mosquito-borne disease. Ecology Letters, 22(10), 1690–1708. 10.1111/ele.13335

Paupy, C., Delatte, H., Bagny, L., Corbel, V., & Fontenille, D. (2009). *Aedes albopictus*, an arbovirus vector: From the darkness to the light. Microbes and Infection, 11(14), 1177–1185. 10.1016/j.micinf.2009.05.005

Petrić, M., Ducheyne, E., Gossner, C. M., Marsboom, C., Nicolas, G., Venail, R., Hendrickx, G., & Schaffner, F. (2021). Seasonality and timing of peak abundance of *Aedes albopictus* in Europe: Implications to public and animal health. Geospatial Health, 16(1), Article 1. 10.4081/gh.2021.996

R Core Team. (2024). R: A Language and Environment for Statistical Computing. R Foundation for Statistical Computing. https://www.R-project.org/

Radici, A., Hammami, P., Cannet, A., L’Ambert, G., Lacour, G., Fournet, F., Garros, C., Guis, H., Fontenille, D., & Caminade, C. (2025). Aedes albopictus is rapidly invading its climatic niche in France: Wider implications for biting nuisance and arbovirus control in Western Europe (p. 2025.02.14.638223). bioRxiv. 10.1101/2025.02.14.638223

Raharimalala, F. N., Ravaomanarivo, L. H., Ravelonandro, P., Rafarasoa, L. S., Zouache, K., Tran-Van, V., Mousson, L., Failloux, A.-B., Hellard, E., Moro, C. V., Ralisoa, B. O., & Mavingui, P. (2012). Biogeography of the two major arbovirus mosquito vectors, Aedes aegypti and Aedes albopictus (Diptera, Culicidae), in Madagascar. Parasites & Vectors, 5(1), 56. 10.1186/1756-3305-5-56

Ribeiro, M. T., Singh, S., & Guestrin, C. (2016). “Why Should I Trust You?”: Explaining the Predictions of Any Classifier. Proceedings of the 22nd ACM SIGKDD International Conference on Knowledge Discovery and Data Mining, 1135–1144. 10.1145/2939672.2939778

Roche, B., Léger, L., L’Ambert, G., Lacour, G., Foussadier, R., Besnard, G., Barré-Cardi, H., Simard, F., & Fontenille, D. (2015). The Spread of Aedes albopictus in Metropolitan France: Contribution of Environmental Drivers and Human Activities and Predictions for a Near Future. PLOS ONE, 10(5), e0125600. 10.1371/journal.pone.0125600

Roiz, D., Boussès, P., Simard, F., Paupy, C., & Fontenille, D. (2015). Autochthonous Chikungunya transmission and extreme climate events in Southern France. PLoS Neglected Tropical Diseases, 9(6), 1–8. 10.1371/journal.pntd.0003854

Roques, L., & Bonnefon, O. (2016). Modelling Population Dynamics in Realistic Landscapes with Linear Elements: A Mechanistic-Statistical Reaction-Diffusion Approach. PLOS ONE, 11(3), e0151217. 10.1371/journal.pone.0151217

Santé Publique France. (2025). Chikungunya, dengue, Zika et West Nile en France hexagonale. Bulletin de la surveillance renforcée du *17 septembre 2025.* https://www.santepubliquefrance.fr/maladies-et-traumatismes/maladies-a-transmission-vectorielle/chikungunya/documents/bulletin-national/chikungunya-dengue-zika-et-west-nile-en-france-hexagonale.-bulletin-de-la-surveillance-renforcee-du-17-septembre-2025

Sherpa, S., Blum, M. G. B., Capblancq, T., Cumer, T., Rioux, D., & Després, L. (2019). Unravelling the invasion history of the Asian tiger mosquito in Europe. Molecular Ecology, 28(9), 2360–2377. 10.1111/mec.15071

Sturiale, S. L., & Armbruster, P. A. (2023). Contrasting effects of an extended fall period and winter heatwaves on the overwintering fitness of diapausing disease vector, Aedes albopictus. Current Research in Insect Science, 4, 100067. 10.1016/j.cris.2023.100067

Székely, G. J., Rizzo, M. L., & Bakirov, N. K. (2007). Measuring and testing dependence by correlation of distances. The Annals of Statistics, 35(6), 2769–2794. 10.1214/009053607000000505

Torina, A., La Russa, F., Blanda, V., Peralbo-Moreno, A., Casades-Martí, L., Di Pasquale, L., Bongiorno, C., Vitale Badaco, V., Toma, L., & Ruiz-Fons, F. (2023). Modelling time-series Aedes albopictus abundance as a forecasting tool in urban environments. Ecological Indicators, 150(April), 110232. 10.1016/j.ecolind.2023.110232

Tran, A., L’Ambert, G., Lacour, G., Benoît, R., Demarchi, M., Cros, M., Cailly, P., Aubry-Kientz, M., Balenghien, T., & Ezanno, P. (2013). A rainfall- and temperature-driven abundance model for Aedes albopictus populations. International Journal of Environmental Research and Public Health, 10(5), 1698–1719. 10.3390/ijerph10051698

Tran, A., Mangeas, M., Demarchi, M., Roux, E., Degenne, P., Haramboure, M., Goff, G. L., Damiens, D., Gouagna, L. C., Herbreteau, V., & Dehecq, J. S. (2020). Complementarity of empirical and processbased approaches to modelling mosquito population dynamics with Aedes albopictus as an example-Application to the development of an operational mapping tool of vector populations. PLoS ONE, 15(1), 1–21. 10.1371/journal.pone.0227407

Urbanski, J., Mogi, M., O’Donnell, D., DeCotiis, M., Toma, T., & Armbruster, P. (2012). Rapid Adaptive Evolution of Photoperiodic Response during Invasion and Range Expansion across a Climatic Gradient. The American Naturalist, 179(4), 490–500. 10.1086/664709

Vanalli, C., Casagrandi, R., Gatto, M., & Bevacqua, D. (2021). Shifts in the thermal niche of fruit trees under climate change: The case of peach cultivation in France. Agricultural and Forest Meteorology, 300(September 2020). 10.1016/j.agrformet.2021.108327

Waldock, J., Chandra, N. L., Lelieveld, J., Proestos, Y., Michael, E., Christophides, G., & Parham, P. E. (2013). The role of environmental variables on Aedes albopictus biology and chikungunya epidemiology. Pathogens and Global Health, 107(5), 224–241. 10.1179/2047773213Y.0000000100

White, S. M., Tegar, S., Purse, B. V., Cobbold, C. A., & Brass, D. P. (2025). Modelling the Lodi, 2023 and Fano 2024, Italy Dengue Outbreaks: The Effects of Control Strategies and Environmental Extremes. Transboundary and Emerging Diseases, 2025(1), 5542740. 10.1155/tbed/5542740

