## Supplementary MM & R for "Temperature is the key weather determinant of *Aedes albopictus* seasonal activity in southern France"

### SI for: Environmental and demographic predictors of *Aedes albopictus* seasonal activity in southern France

#### Additional materials

##### Metelmann model parameters

In the modelling framework established by Metelmann et al., (2019) both the carrying capability $K_{it}$ and the hatching $h_{it}$ of a site $i$ at time $t$ depend on host density $H_{it}$ and rainfall $r_{it}$:

$K_{it}=\lambda\frac{1- \alpha_{evap}}{1- {\alpha_{evap}}^{t}}\sum_{x=1}^{t} {\alpha_{evap}}^{\left( t-x \right)}( \alpha_{rain}r_{ix}+ \alpha_{dens}H_{ix}$)

$$h_{it}=\left( 1- \varepsilon_{rat} \right)\frac{\left( 1+\varepsilon_{0} \right)e^{- \varepsilon_{var}\left( r_{it}-\varepsilon_{opt} \right)^{2}}}{e^{- \varepsilon_{var}\left( r_{it}-\varepsilon_{opt} \right)^{2}}+ \varepsilon_{0}}+ \varepsilon_{rat}\frac{\varepsilon_{dens}}{\varepsilon_{dens}+ e^{-\varepsilon_{fac}H_{it}}}$$

Note that, in the main text, we have indicated the physical quantity of each environmental driver, without distinguishing among different functional forms. For example, both adult mortality
$\mu_{A}(\bar{T})$ and survival probability of diapausing eggs $\gamma(T_{\min\left( DJF \right)})$ depends on temperature, but the former refers to its daily average, while the latter depends on the minimum value in the previous winter.

Here we present the table of the values of the parameters of the abovementioned equations and those of the compartmental model presented in the main text, with the corresponding source:

Table SI1 – Metelmann model parameters

| **Parameter** | **Meaning** | **Value/formula** | **Source** |
| --- | --- | --- | --- |
| ${CTT}_{S}$ | critical temperature over one week in spring (°C ) | $11.0$ | (Metelmann et al., 2019) |
| ${CPP}_{S}$ | critical photoperiod in spring (hours) | $11.25$ | (Metelmann et al., 2019) |
| $\sigma(T,P)$ | spring hatching rate (day^-1^) | $\left\{ \begin{aligned} 0 if T_{7}<\mathrm{CTT}_{S} or P< \mathrm{CPP}_{S} \\ 0.1 otherwise \end{aligned} \right.$ | (Metelmann et al., 2019) |
| ${CPP}_{A}(L)$ | critical photoperiod in autumn (hours) | $10.058+0.08965L$ | (Metelmann et al., 2019) |
| $\omega(P)$ | fraction of eggs going into diapause | $\left\{ \begin{aligned} 0 if P<\mathrm{CPP}_{A} or day< 183 \\ 0.5 otherwise \end{aligned} \right.$ | (Metelmann et al., 2019) |
| $\delta_{E}$ | normal egg development rate (day^-1^) | $1/7.1$ | (Metelmann et al., 2019) |
| $\delta_{J}(T)$ | Juvenile development rate (day^-1^) | $1/(83.85- 4.89 T+ 0.08 T^{2})$ | (Metelmann et al., 2019) |
| $\delta_{I}(T)$ | first pre-blood meal rate (day^-1^) | $1/(50.1- 3.574 T+ 0.069 T^{2})$ | (Metelmann et al., 2019) |
| $\mu_{E}$ | egg mortality rate (day^-1^) | $-ln(0.955 e^{-0.5{(\frac{T-18.8}{21.53})}^{6}} )$ | (Metelmann et al., 2019) |
| $\mu_{J}$ | juvenile mortality rate (day^-1^) | $-ln(0.977 e^{-0.5{(\frac{T-21.8}{16.6})}^{6}} )$ | (Metelmann et al., 2019) |
| $\mu_{A}(\bar{T})$ | adult mortality rate (day^-1^) | $-ln(0.677 e^{-0.5{(\frac{\bar{T}-20.29}{13.2})}^{6}}0.069 \bar{T}^{0.1} )$ | (Metelmann et al., 2019) |
| $\gamma(T_{\min\left( DJF \right)})$ | survival probability of diapausing eggs (winter^-1^) | $0.93 e^{-0.5{(\frac{T_{\min\left( \mathrm{DJF} \right)}-11.68}{15.67})}^{6}}$ | (Metelmann et al., 2019) |
| $\beta(T)$ | egg laying rate (day^-1^) | $\left\{ \begin{aligned} 33.2 e^{-0.5{(\frac{T - 70.3}{14.1})}^{2}}{(38.8-T)}^{1.5} if T\leq38.8 \\ 0 otherwise \end{aligned} \right.$ | (Metelmann et al., 2019) |
| $\lambda$ | capacity parameter (larvae days ha^-1^) | ${10}^{6}$ | (Metelmann et al., 2019) |
| $\alpha_{evap}$ | Normalization parameter evaporation | $0.9$ | (Metelmann et al., 2019) |
| $\alpha_{dens}$ | Normalization parameter of host density (km²) | ${10}^{-5}$ | (Metelmann et al., 2019) |
| $\alpha_{rain}$ | Normalization parameter of rain (mm^-2^) | ${10}^{-2}$ | (Metelmann et al., 2019) |
| $\varepsilon_{rat}$ | Proportion of hatching due to rain | $0.2$ | (Metelmann et al., 2019) |
| $\varepsilon_{0}$ | Correcting additive parameter for rain-related hatching | $1.5$ | (Metelmann et al., 2019) |
| $\varepsilon_{var}$ | Stretching parameter for rain-related hatching (mm^-2^) | $0.05$ | (Metelmann et al., 2019) |
| $\varepsilon_{opt}$ | Correcting additive parameter for human-induced hatching (mm) | $8$ | (Metelmann et al., 2019) |
| $\varepsilon_{dens}$ | for rain-related hatching | ${10}^{-2}$ | (Metelmann et al., 2019) |
| $\varepsilon_{fac}$ | Stretching parameter for human-induced hatching (km^-2^) | ${10}^{-2}$ | (Metelmann et al., 2019) |

The model schemes are represented here:

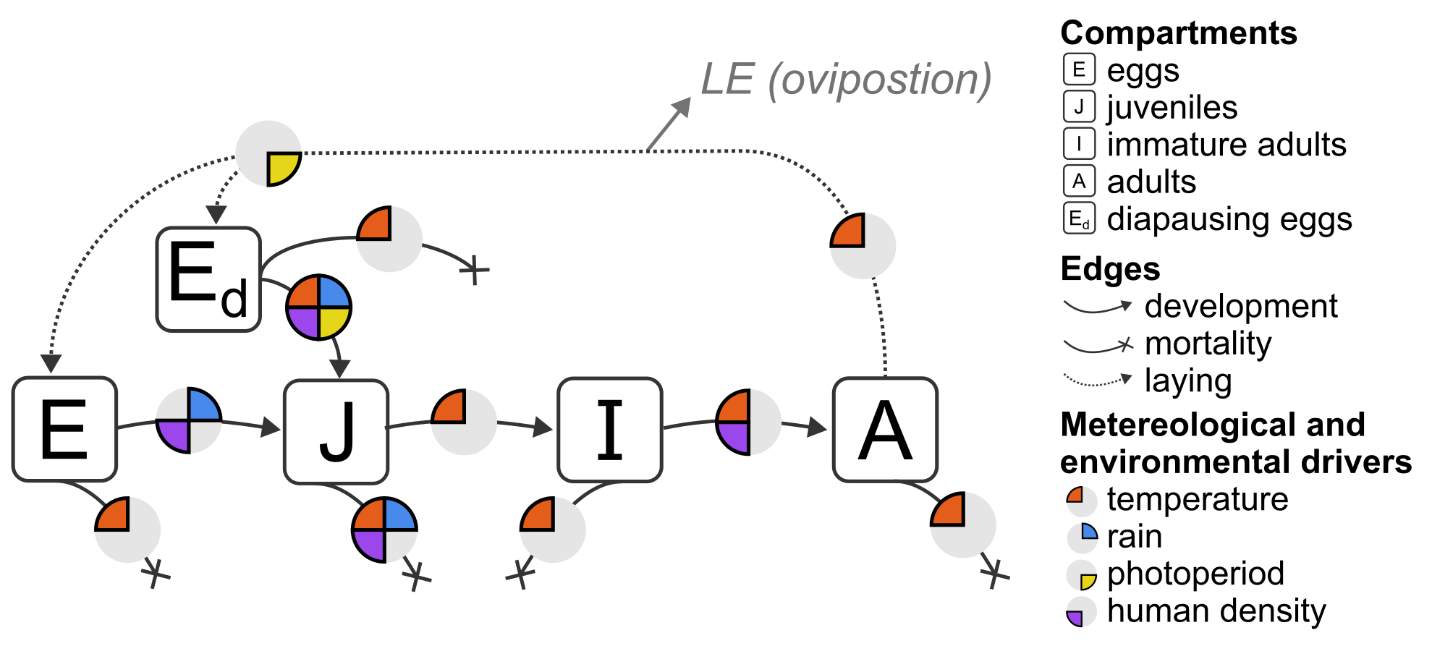

Figure SI1 - Layout of the weather-driven Metelmann model describing Ae. albopictus dynamics, edited from (Radici et al., 2025)

##### Physical quantities in models

In the following table, we report the explicit definition of the weather and environmental variables:

Table SI2– Weather variables and specifications

| **Weather and environmental variable** | **Meaning** | **Model** |
| --- | --- | --- |
| $T$ | Instantaneous temperature (°C) | Metelmann |
| $\bar{T}$ | Mean daily temperature (°C) | Metelmann |
| $T_{7}$ | Mean temperature of the last week (°C) | Metelmann |
| $T_{\min\left( DJF \right)}$ | Lowest winter (December-January-February) temperature (°C) | Metelmann |
| $R$ | Daily rainfall (mm) | Metelmann |
| $H$ | (Human) Host density (hab/km^2^) | Metelmann |
| $P$ | Photoperiod (in hour) | Metelmann |
| $L$ | Latitude (°) | Metelmann |

##### Ovitrap collection details

In the following table, details about the ovitrap records samplings. In rare cases in which the average temperature was missing, it was computed as the mean of the maximal and minimal daily temperature.

Table SI3 – Summary about ovitrap and weather recording per site

| **Site** | **Mean number of active ovitraps per week** | **Series length** | **Years** | **Corresponding Météo France weather station** | **Human density (km^-2^, average from GPW)** |
| --- | --- | --- | --- | --- | --- |
| Pérols | 19.1 | 64 | 2023-2024 | Montpellier-aéroport | 1013.7 |
| Murviels-les-Montpellier | 27.7 | 43 | 2023-2024 | Montarnaud | 102.1 |
| Saint-Médard-en-Jalle | 19.9 | 47 | 2023-2024 | Bordeaux-Merignac | 991.4 |
| Bayonne | 20.5 | 41 | 2023-2024 | Biarritz-Pays-Basque | 826.0 |

Table SI4 - Extended summary of ovitrap and weather data per site, year and season

| Site | Year | Season | Weather | | Entomology | | |
| --- | --- | --- | --- | --- | --- | --- | --- |
|  |  |  | Average T (°C) | Cumulated rainfall in season (mm) | Mean abundance (egg/ovitrap) | Begin of activity (week) | End of activity (week) |
| Pérols | 2023 | Spring | 21.1 | 85.4 | 19.6 | 19 | 46 |
|  |  | Summer | 25.2 | 15.9 | 30.7 |  |  |
|  |  | Autumn | 20.3 | 54.3 | 43.7 |  |  |
|  |  | Nov-> Ap | 10.6 | 105.7 | 0.3 |  |  |
|  | 2024 | Spring | 19.4 | 105.4 | 14.3 | 20 | 51 |
|  |  | Summer | 25.6 | 26.6 | 38.5 |  |  |
|  |  | Autumn | 18.7 | 118.1 | 36.0 |  |  |
|  |  | Nov-> Ap | 11.0 | 329.6 | 0.8 |  |  |
| Mur.-les-Montpellier | 2023 | Spring | 20.0 | 153.6 | 23.7 | 18 | 48 |
|  |  | Summer | 24.9 | 13.6 | 40.9 |  |  |
|  |  | Autumn | 18.9 | 159.3 | 28.8 |  |  |
|  |  | Nov-> Ap | 9.6 | 120.8 | 0.5 |  |  |
|  | 2024 | Spring | 18.5 | 140.4 | 8.3 | 19 | 50 |
|  |  | Summer | 25.1 | 29.4 | 28.4 |  |  |
|  |  | Autumn | 17.3 | 234.1 | 16.7 |  |  |
|  |  | Nov-> Ap | 10.0 | 428.2 | 0.3 |  |  |
| Bayonne | 2023 | Spring | 18.7 | 161.2 | 6.0 | 19 | - |
|  |  | Summer | 21.4 | 148.1 | - |  |  |
|  |  | Autumn | 20.1 | 355.2 | - |  |  |
|  |  | Nov-> Ap | 11.1 | 1,072.8 | 0.0 |  |  |
|  | 2024 | Spring | 16.7 | 387.8 | 3.5 | 17 | 51 |
|  |  | Summer | 21.3 | 70.2 | 22.8 |  |  |
|  |  | Autumn | 17.3 | 536.7 | 20.2 |  |  |
|  |  | Nov-> Ap | 11.7 | 824.7 | 0.4 |  |  |
| St.-Médard-en-Jalle | 2023 | Spring | 19.8 | 156.0 | 8.2 | 19 | - |
|  |  | Summer | 22.0 | 55.6 | - |  |  |
|  |  | Autumn | 19.4 | 280.9 | - |  |  |
|  |  | Nov-> Ap | 9.8 | 693.9 | 0.0 |  |  |
|  | 2024 | Spring | 17.4 | 219.6 | 3.6 | 20 | 49 |
|  |  | Summer | 22.0 | 60.9 | 32.2 |  |  |
|  |  | Autumn | 16.4 | 206.7 | 16.4 |  |  |
|  |  | Nov-> Ap | 10.3 | 529.6 | 0.1 |  |  |

##### Cross correlation maps

In the following figures, we show the cross correlation maps between lagged environmental drivers and ovitrap measurements. In Figure SI2, we show the correlation for the presence-absence model, while in Figure SI3 for the continuous abundance model.

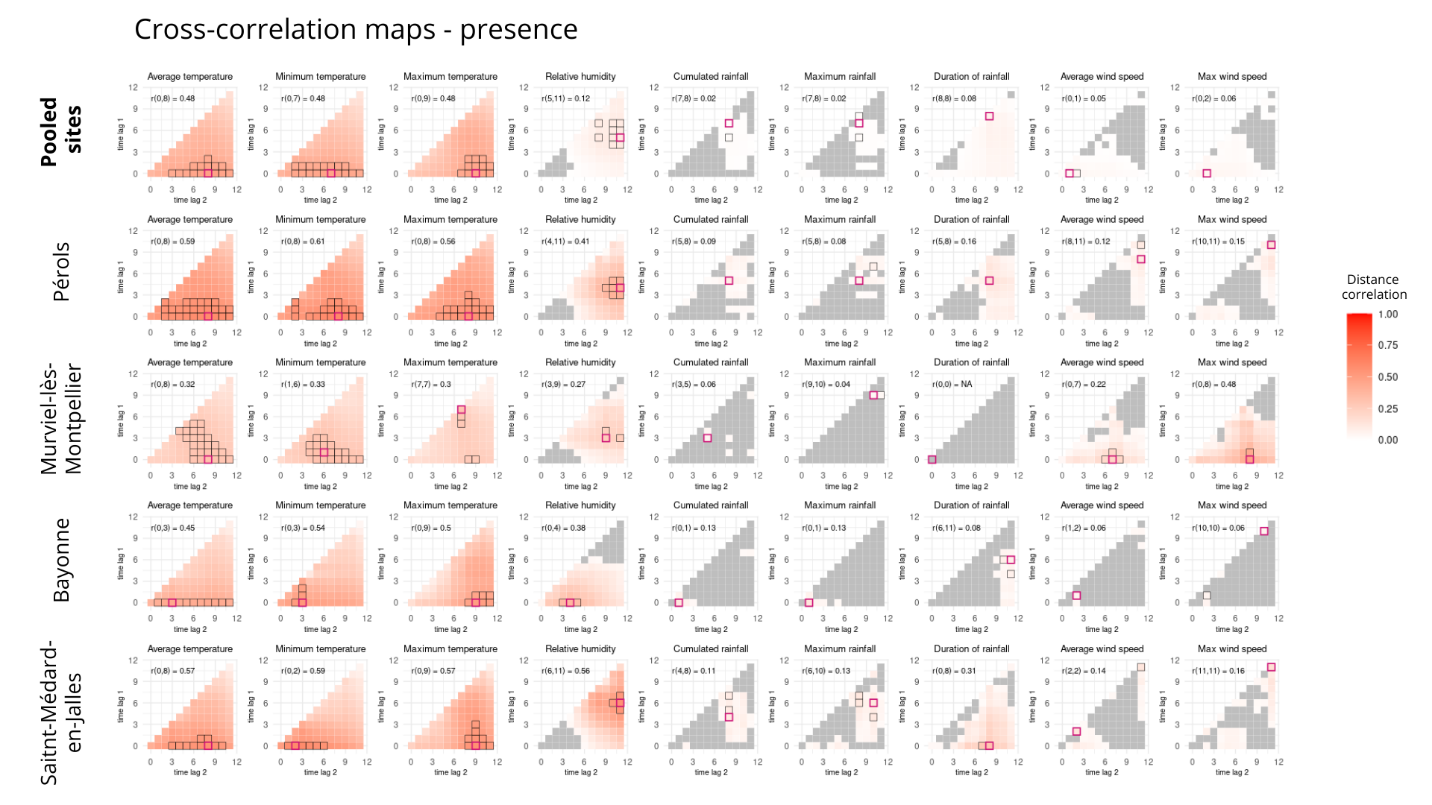

Figure SI2 - Cross correlation maps for each site (rows) and environmental variables (columns) against ovitrap observations of presence and absence.

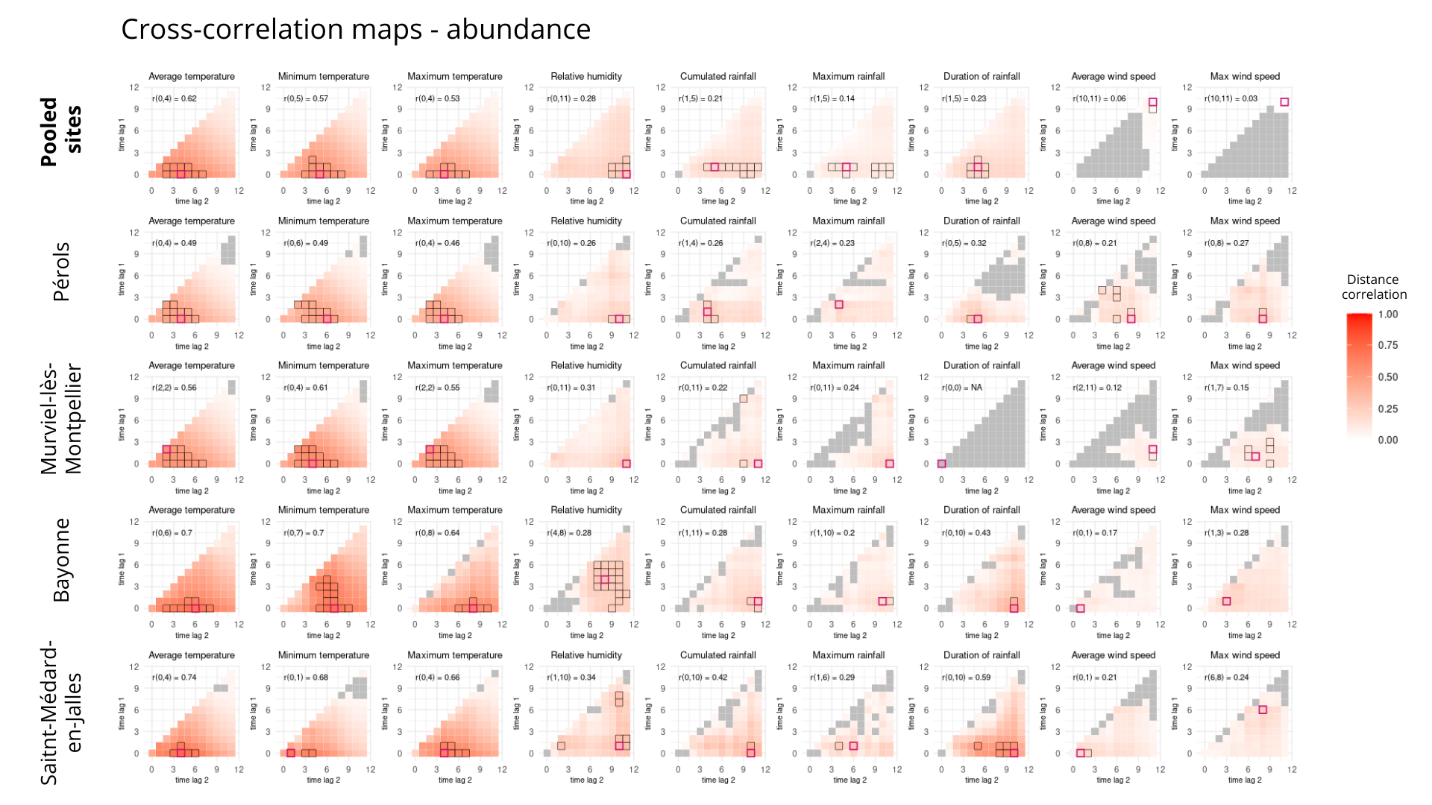

Figure SI3 - Cross correlation maps for each site (rows) and environmental variables (columns) against ovitrap observations of oviposition abundance.

##### LIME site by site

In the following, we report the detailed LIME (Local Interpretable Model-agnostic Explanations) for each site: Pérols (Figure SI4), Murviels les Montpellier (Figure SI5), Bayonne (Figure SI6), Saint Médard-en-Jalle (Figure SI7).

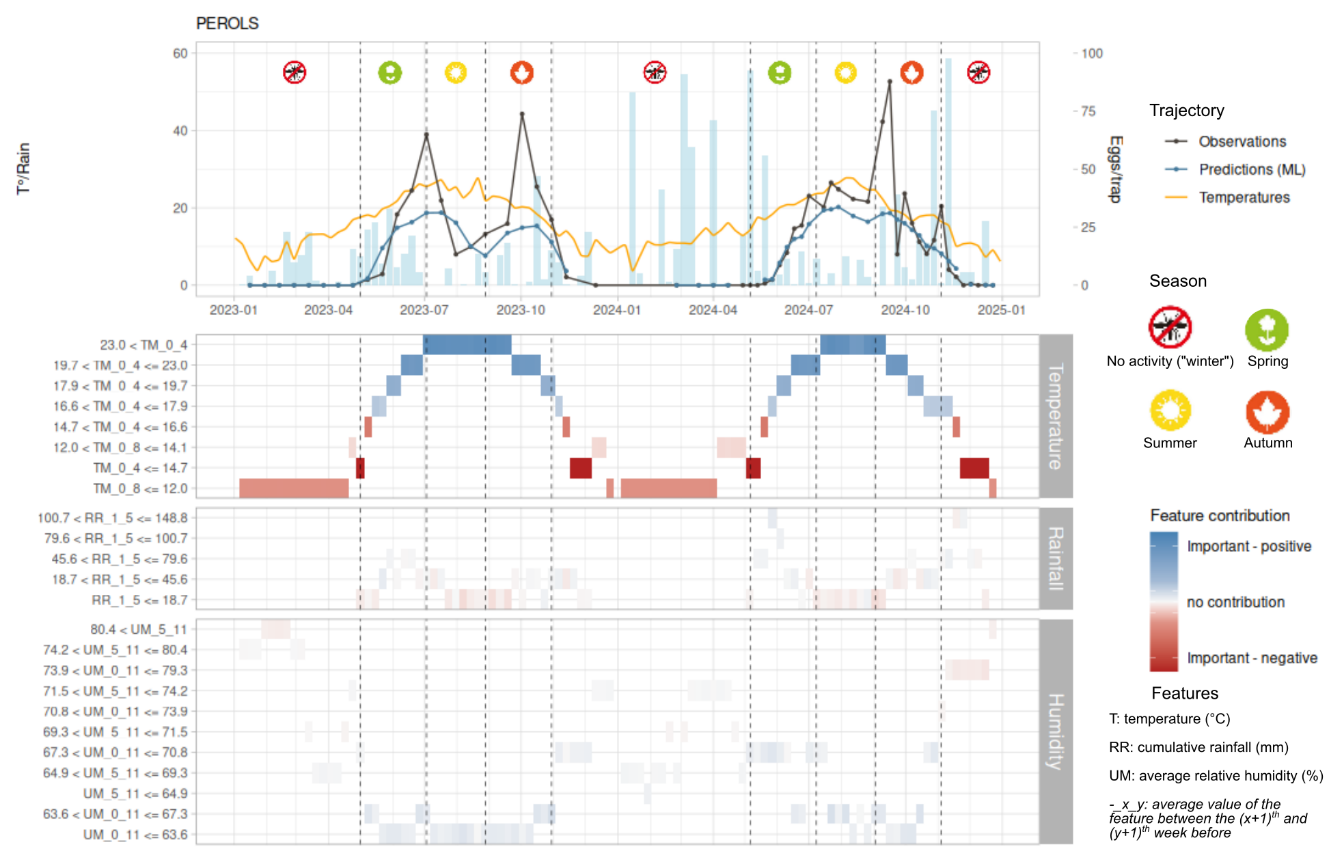

Figure SI4 - Local contribution of environmental predictors to predicted mosquito abundance in Pérols using LIME

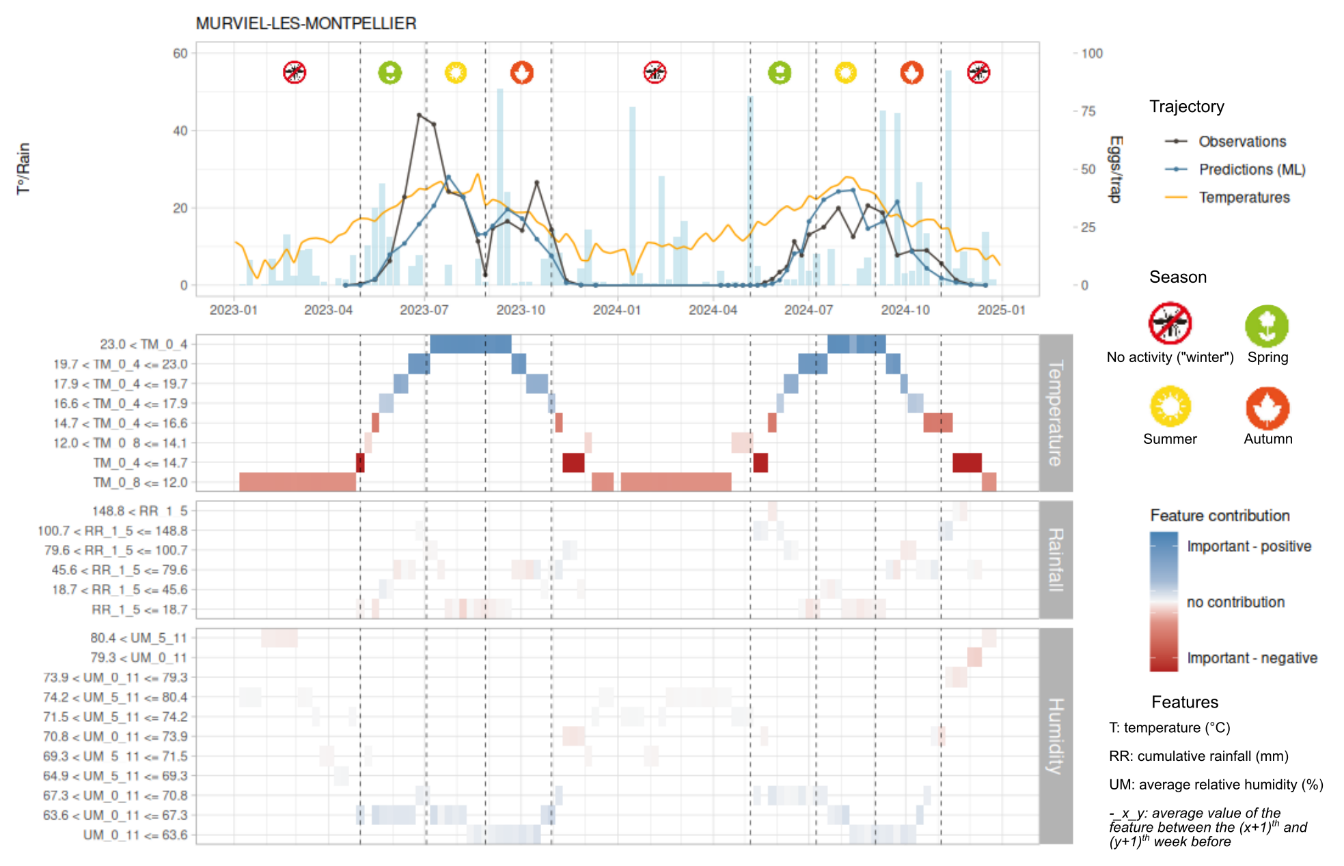

Figure SI5 - Local contribution of environmental predictors to predicted mosquito abundance in Murviels-les-Montpellier using LIME

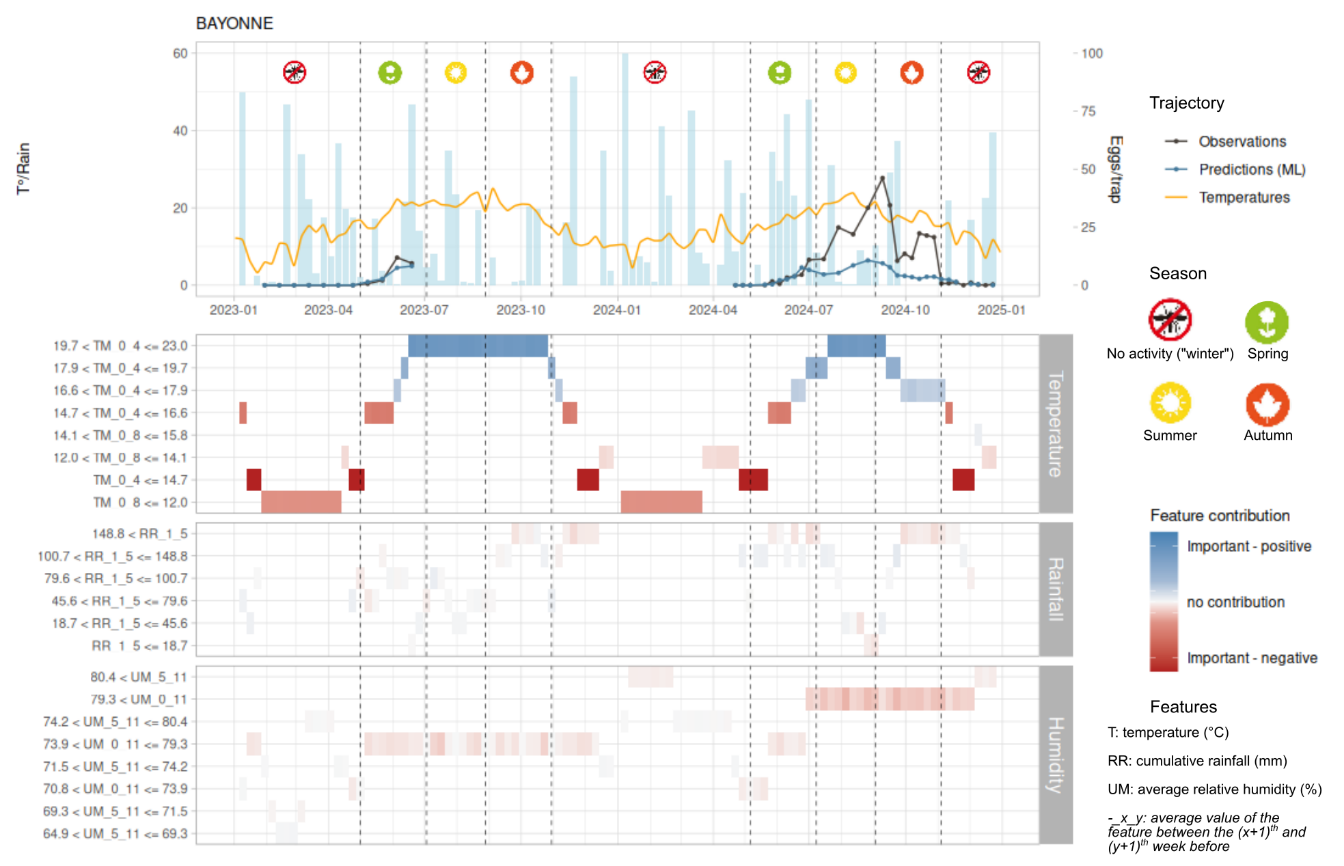

Figure SI6 - Local contribution of environmental predictors to predicted mosquito abundance in Bayonne using LIME

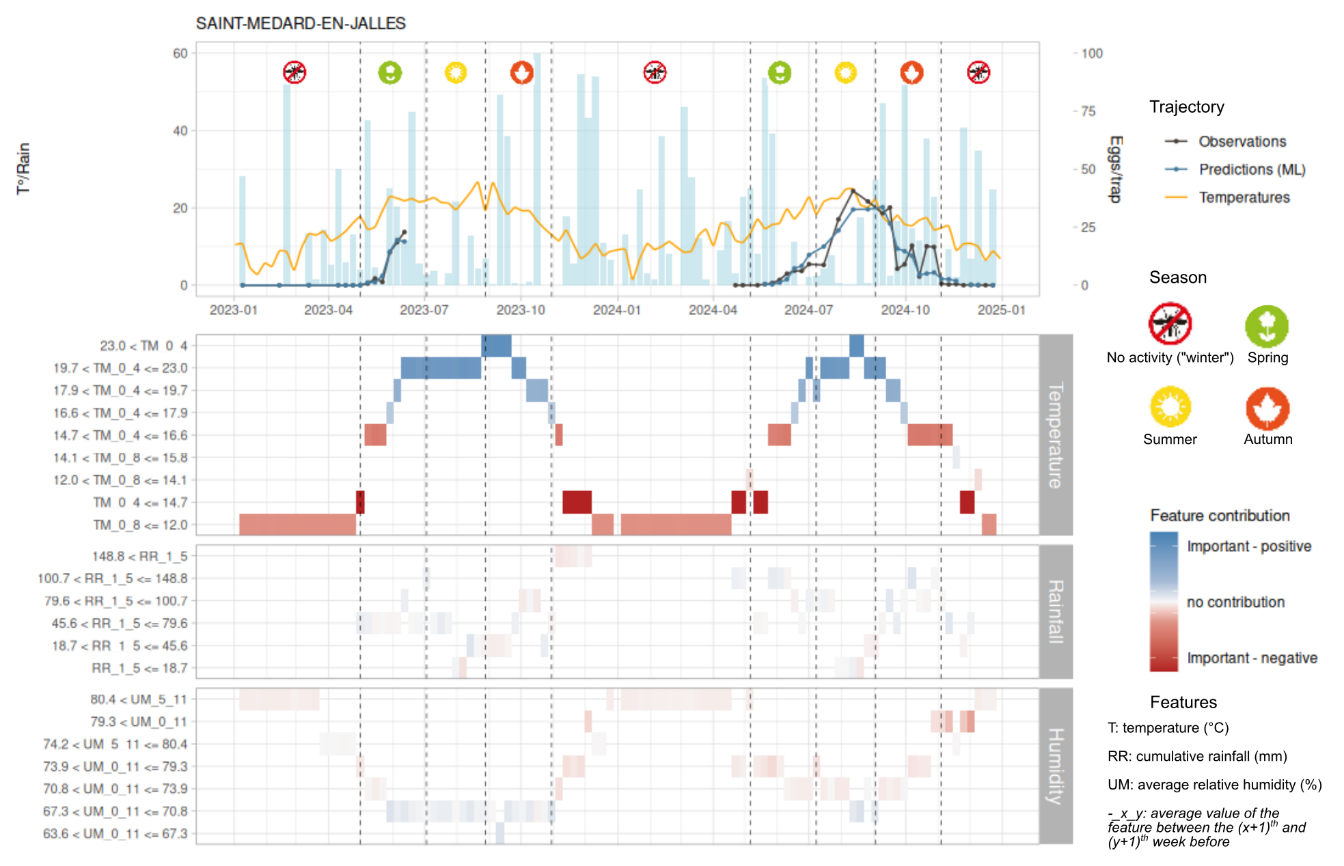

Figure SI7 - Local contribution of environmental predictors to predicted mosquito abundance in Saint Médard en Jalle using LIME

#### Additional references

Metelmann, S., Caminade, C., Jones, A.E., Medlock, J.M., Baylis, M., Morse, A.P., 2019. The UK’s suitability for Aedes albopictus in current and future climates. Journal of the Royal Society Interface 16. https://doi.org/10.1098/rsif.2018.0761

Radici, A., Hammami, P., Cannet, A., L’Ambert, G., Lacour, G., Fournet, F., Garros, C., Guis, H., Fontenille, D., Caminade, C., 2025. Aedes albopictus is rapidly invading its climatic niche in France: wider implications for biting nuisance and arbovirus control in Western Europe. https://doi.org/10.1101/2025.02.14.638223
